# Structural robustness affects the engineerability of aminoacyl-tRNA synthetases for genetic code expansion

**DOI:** 10.1101/829028

**Authors:** Katherine T. Grasso, Megan Jin Rae Yeo, Christen M. Hillenbrand, Elise D. Ficaretta, James S. Italia, Rachel L. Huang, Abhishek Chatterjee

## Abstract

The ability to engineer the substrate specificity of natural aminoacyl-tRNA synthetase/tRNA pairs facilitates the site-specific incorporation of noncanonical amino acids (ncAAs) into proteins. The *Methanocaldococcus jannaschii* derived tyrosyl-tRNA synthetase (MjTyrRS)/tRNA pair has been engineered to incorporate numerous ncAAs into protein expressed in bacteria. However, it cannot be used in eukaryotic cells due to cross-reactivity with its host counterparts. The *E. coli* derived tyrosyl-tRNA synthetase (EcTyrRS)/tRNA pair offers a suitable alternative to this end, but a much smaller subset of ncAAs has been genetically encoded using this pair. Here we report that this discrepancy, at least partly, stems from the lower structural robustness of EcTyrRS relative to MjTyrRS. We show that engineered TyrRS mutants in general exhibit significantly lower thermostability relative to their wild-type counterparts. Derived from a thermophilic archaeon, MjTyrRS is a remarkably sturdy protein and tolerates extensive active site engineering without a catastrophic loss of stability at physiological temperature. In contrast, EcTyrRS exhibits significantly lower thermostability, rendering some of its engineered mutants insufficiently stable at physiological temperature. Our observations identify the structural robustness of an aaRS as an important factor that significantly influences how extensively it can be engineered. To overcome this limitation, we have further developed chimeras between EcTyrRS and its homolog from a thermophilic bacteria, which offer an optimal balance between thermostability and activity. We show that the chimeric bacterial TyrRSs show enhanced tolerance for destabilizing active site mutations, providing a potentially more engineerable platform for genetic code expansion.

---

Noncanonical amino acid (ncAA) mutagenesis of proteins in living cells has emerged as a powerful technology with enormous potential.^*1–5*^ A ncAA of interest can be co-translationally incorporated using an orthogonal aminoacyl-tRNA synthetase (aaRS)/tRNA pair in response to a nonsense or frameshift codon.^*1–5*^ Central to this technology is the ability to engineer the substrate specificity of a natural aaRS through directed evolution. Many useful ncAAs have been genetically encoded in *E. coli* using the *Methanocaldococcus jannaschii* derived tyrosyl-tRNA synthetase (MjTyrRS)/tRNA pair, including those containing bioconjugation handles, photo-affinity probes, biophysical probes, models for natural post-translational modification, etc.^*1, 2, 5*^ While some of these functionalities can also be genetically encoded using other aaRS/tRNA pairs, several others (e.g., those modeling natural post-translational modifications) are reliant on the unique architecture of the TyrRS active site.^*1, 2, 5*^ Unfortunately, however, this enabling toolset cannot be used in eukaryotic cells, as the archaea-derived MjTyrRS/tRNA pair cross-reacts with its eukaryotic counterpart. Typically, bacteria-derived aaRS/tRNA pairs are suitable for ncAA incorporation in eukaryotes, as they tend to be orthogonal in these cells.^*1, 3, 5–7*^ Indeed, the *E. coli* derived tyrosyl-tRNA synthetase (EcTyrRS)/tRNA pair has been established for ncAA incorporation in eukaryotic cells.^*8–12*^ It was first engineered to incorporate ncAAs into proteins expressed in eukaryotic cells nearly two decades ago. Yet, the ncAA-toolbox developed using this pair remains surprisingly limited, particularly when compared to the remarkable success of the MjTyrRS/tRNA pair during the same time period.^*1–3, 5, 13*^ The ability to recapitulate the success of the MjTyrRS/tRNA platform using a bacterial TyrRS/tRNA pair will significantly expand the scope of the genetic code expansion (GCE) technology in eukaryotes by providing access to structurally unique ncAAs that are challenging to genetically encode using alternative aaRS/tRNA pairs.

The limited success of the EcTyrRS/tRNA pair can be, at least partially, attributed to the challenges associated with the directed evolution platform used to alter its substrate specificity.^*1, 3, 7, 13*^ Unlike MjTyrRS, which can be readily engineered using a facile *E. coli* based directed evolution system, a more cumbersome yeast-based selection scheme is needed to engineer EcTyrRS.^*8*^ To address this challenge, we recently developed a novel strategy that involves the development of unique *E. coli* strains (ATMY strains), where the endogenous EcTyrRS/tRNA pair is functionally substituted with an archaeal counterpart.^*1, 3, 7, 13*^ We have demonstrated that such strains can be generated without incurring a significant growth penalty.^*13*^ The ‘liberated’ EcTyrRS/tRNA pair can be subsequently established in the resulting ATMY strains as an orthogonal nonsense suppressor. This has enabled the use of the facile *E. coli* based directed evolution platform to engineer the substrate specificity of EcTyrRS.^*13*^

Although the ability to rapidly engineer the EcTyrRS/tRNA pair using our facile directed evolution platform has provided access to several new ncAAs, in some instances, the resulting engineered mutants demonstrated poor activity. For example, we attempted to develop an EcTyrRS mutant that efficiently charges p-benzoylphenylalanine (pBpA), a powerful photoaffinity probe that has been useful for capturing weak and transient molecular interactions.^*8, 14–17*^ Even though EcTyrRS was previously engineered using the yeast-based selection platform to selectively charge pBpA, the utility of this mutant has been limited due to its weak activity. When we subjected a large EcTyrRS active site mutant library to our facile selection system, we identified the same mutant that was previously developed by selection in yeast. This indicated that the observed set of mutations indeed optimally recodes the EcTyrRS active site for charging pBpA. The activity of this pBpA-selective EcTyrRS was somewhat low, as measured in the ATMY *E. coli* strain using the sfGFP-151-TAG reporter (Figure S1). A systematic investigation identified that the EcTyrRS-pBpA mutant was largely insoluble in *E. coli*, potentially explaining its poor activity (Figure 1B). Western blot analysis of the soluble and insoluble fractions of cell-free extracts of *E. coli* expressing polyhistidine-tagged EcTyrRS-WT and EcTyrRS-pBpA revealed that the latter is nearly exclusively found in the insoluble fraction (Figure 1B). In contrast, a large portion of the wild-type EcTyrRS was found in the soluble fraction.

**Figure 1.**
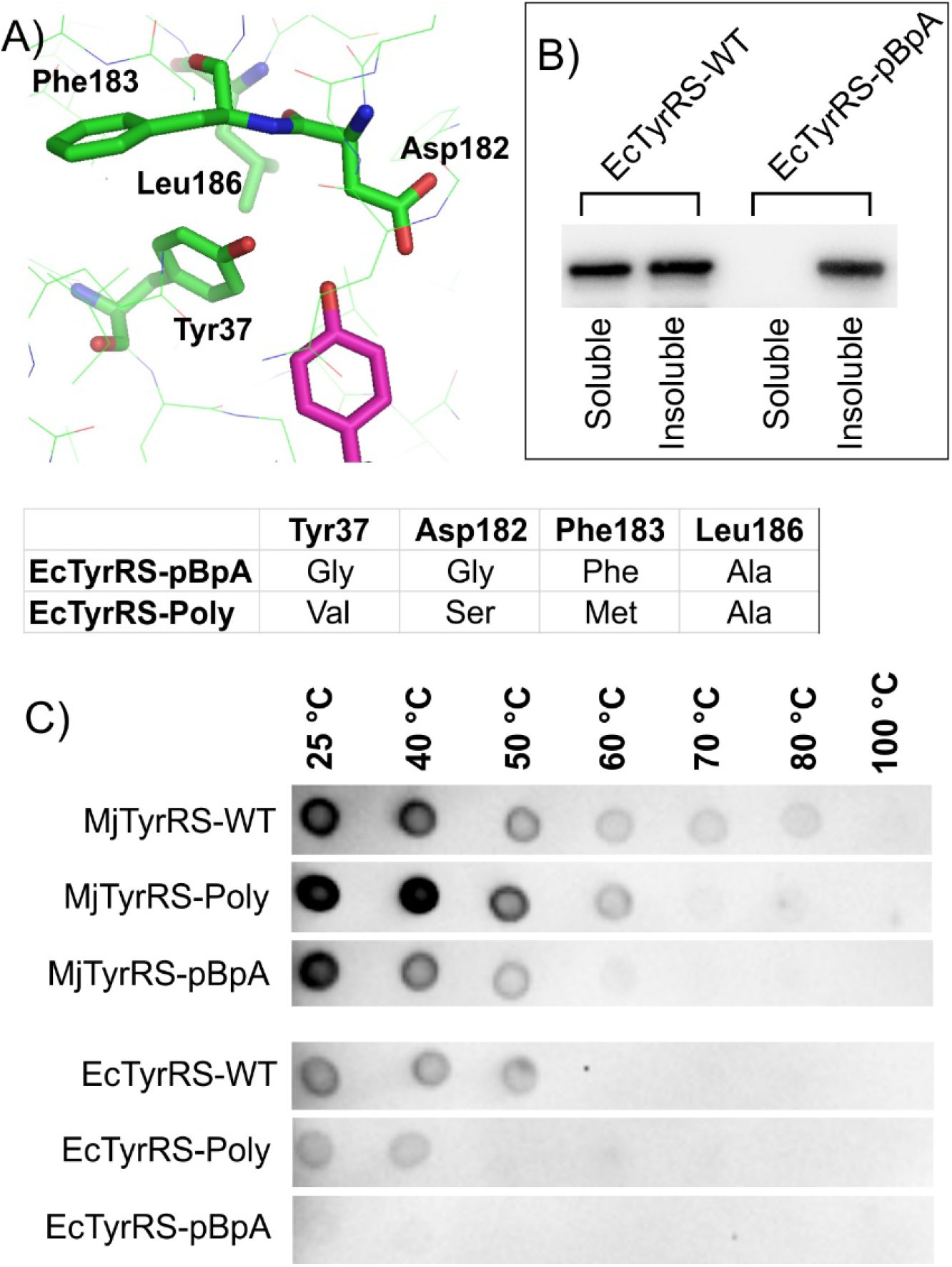
Engineered TyrRS mutants exhibit lower thermostability. A) EcTyrRS active site, showing bound tyrosine in magenta, and highlighting mutated residues in engineered variants (shown below). B) Western blot analysis of soluble and insoluble fractions of *E. coli* cell free extract expressing EcTyrRS-WT and EcTyrRS-pBpA reveals that the latter is largely insoluble. C) Thermal shift assay of various TyrRS variants. Dot-blot was performed (using an anti-polyhistidine antibody) on the soluble fraction of *E. coli* cell-free extracts expressing the indicated TyrRS variants (left), after incubation at the indicated temperature (top).

It has been previously observed that when a protein is subjected to directed evolution to attain an altered function, the stability of the resulting mutants is often compromised.^*18–21*^ Consequently, the extent to which a protein can be engineered is often limited by how stable it is. We hypothesized that the success in engineering MjTyrRS, an enzyme derived from a thermophilic archaeon, is likely facilitated by its high structural stability. In contrast, EcTyrRS, derived from a mesophilic bacteria, may be a less stable scaffold and have a lower tolerance for active site mutations. To test this notion, we took advantage of a modified cellular thermal shift assay (CETSA).^*22, 23*^ In this assay, cell-free extract expressing a target protein is heated to increasing temperatures, and the amount of remaining protein in the soluble fraction is subsequently tested by immunoblotting. The temperature range at which a protein is lost from the soluble fraction provides an estimate of its thermostability. In addition to EcTyrRS-WT and EcTyrRS-pBpA, we also tested a polyspecific EcTyrRS mutant (EcTyrRS-Poly) that is highly active (Figure 1A). For comparison, we included MjTyrRS-WT, as well as two of its comparable engineered mutants: one selective for pBpA (MjTyrRS-pBpA),^*24*^ and another that exhibits ncAA polyspecificity (MjTyrRS-Poly) (Figure S2).^*25*^ All of these proteins encoded an N-terminal hexahystidine tag to facilitate their detection in a dot-blot assay using an anti-polyhistidine antibody. As expected, MjTyrRS was found to be highly thermostable, maintaining solubility up to 80 °C (Figure 1C). In contrast, EcTyrRS was much less stable, and was lost from the soluble fraction between 50 °C and 60 °C (Figure 1C). These values are consistent with previously reported thermostability measurements.^*26*^ All of the engineered mutants exhibited reduced stability relative to their wild-type counterparts. MjTyrRS-pBpA mutant was slightly less stable than its polyspecific counterpart, but both were soluble at physiological temperature. In contrast, for EcTyrRS, only the polyspecific mutant was soluble at physiological temperature; the pBpA selective mutant was not detected in the soluble fraction even at the lowest temperature tested (Figure 1C). These observations support the hypothesis that the lower stability of EcTyrRS negatively impacts its engineerability. While some of its active site mutants, such as EcTyrRS-Poly, are adequately stable and active under physiological conditions, more destabilizing mutants such as EcTyrRS-pBpA are not viable.

To overcome this challenge, we considered the possibility of adapting a TyrRS from a thermophilic bacterium, which might offer a higher degree of engineerability relative to EcTyrRS. Several aminoacyl-tRNA synthetases derived from the thermophilic bacterium *Geobacillus stearothermophilus* have been purified and structurally characterized.^*27–29*^ TyrRS from this bacterium (GsTyrRS) is homologous to EcTyrRS (Figure S3) but is significantly more thermostable, offering an attractive scaffold for engineering ncAA-selective mutants.^*26*^ We tested the stability and the activity of GsTyrRS, using the CETSA and the sfGFP-151-TAG expression assay, as described above (Figure 2B). GsTyrRS was indeed significantly more stable than EcTyrRS, but its activity was somewhat lower in *E. coli* (Figure 2B, 2C). This is not surprising; enzymes derived from thermophilic bacteria often exhibit weaker activity at lower temperature.^*30*^ We thought that it may be possible to create a chimera from EcTyrRS and GsTyrRS that exhibit an optimal balance of stability and activity. Indeed, it has been previously shown that such chimeric enzymes can be excellent scaffolds for protein engineering.^*31, 32*^ We created and tested several chimeras between EcTyrRS and GsTyrRS and identified two, Ch2TyrRS and Ch6TyrRS (Figure 2A), which show higher levels of activity (Figure 2C) and intermediate thermostability (Figure 2B) relative to their wild-type progenitors, when expressed in ATMY *E. coli*. Next, we generated and tested pBpA-selective mutants of these TyrRS variants. Both Ch2TyrRS-pBpA and Ch6TyrRS-pBpA were more thermostable than EcTyrRS-pBpA, and were soluble at physiological temperature (Figure 3A). This was reflected by the significantly higher degree of activity observed for these enzymes (Figure 3B). Interestingly, even though GsTyrRS was the most thermostable, the corresponding pBpA-selective mutant was found to be largely insoluble and poorly active when expressed in *E. coli* (Figure 3A, 3B). While the basis of this observation is unclear, it indicates that the chimeras may be better suited for engineering new ncAA-selective variants.

**Figure 2.**
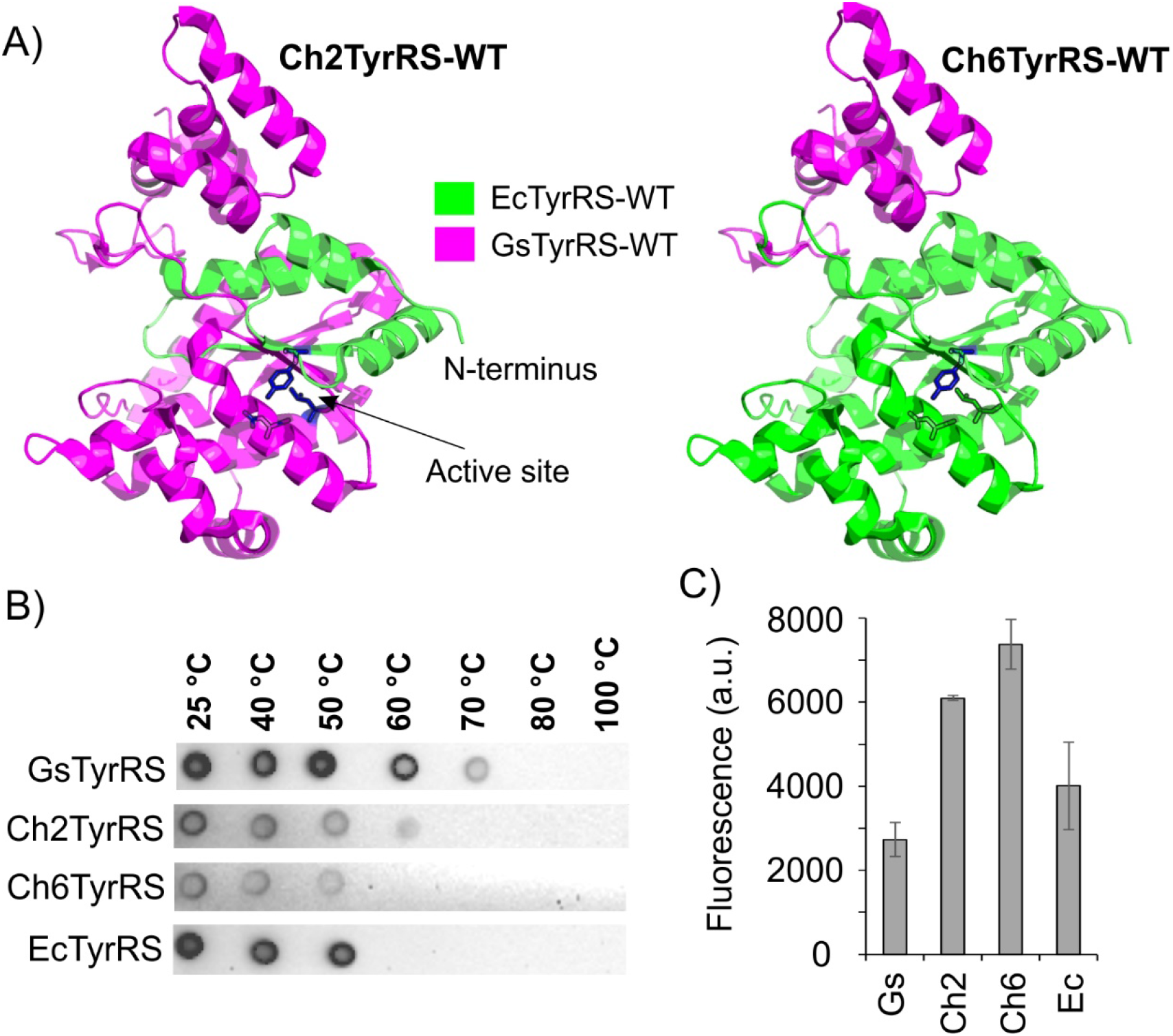
Bacterial TyrRS chimeras created from GsTyrRS and EcTyrRS exhibit higher activity and intermediate thermostability. A) Two chimeras, Ch2TyrRS and Ch6TyrRS, created by fusing EcTyrRS (green) and GsTyrRS (magenta) sequences. The EcTyrRS crystal structure was used to highlight the progenitor sequences in the two chimeras. B) Thermal shift assay of the two chimeras, as well as their wild-type progenitors, in *E. coli* cell-free extract. Dot-blot was performed (using an anti-polyhistidine antibody) on the soluble fraction of *E. coli* cell-free extracts expressing the indicated TyrRS variants (left), after incubation at the indicated temperature (top). C) Activity of the TyrRS variants in ATMY *E. coli* strains, measured using the expression of sfGFP-151TAG reporter in the presence of tRNA_CUA_^Tyr^. Characteristic fluorescence of the full-length sfGFP-151-TAG reporter was measured in resuspended cells.

**Figure 3.**
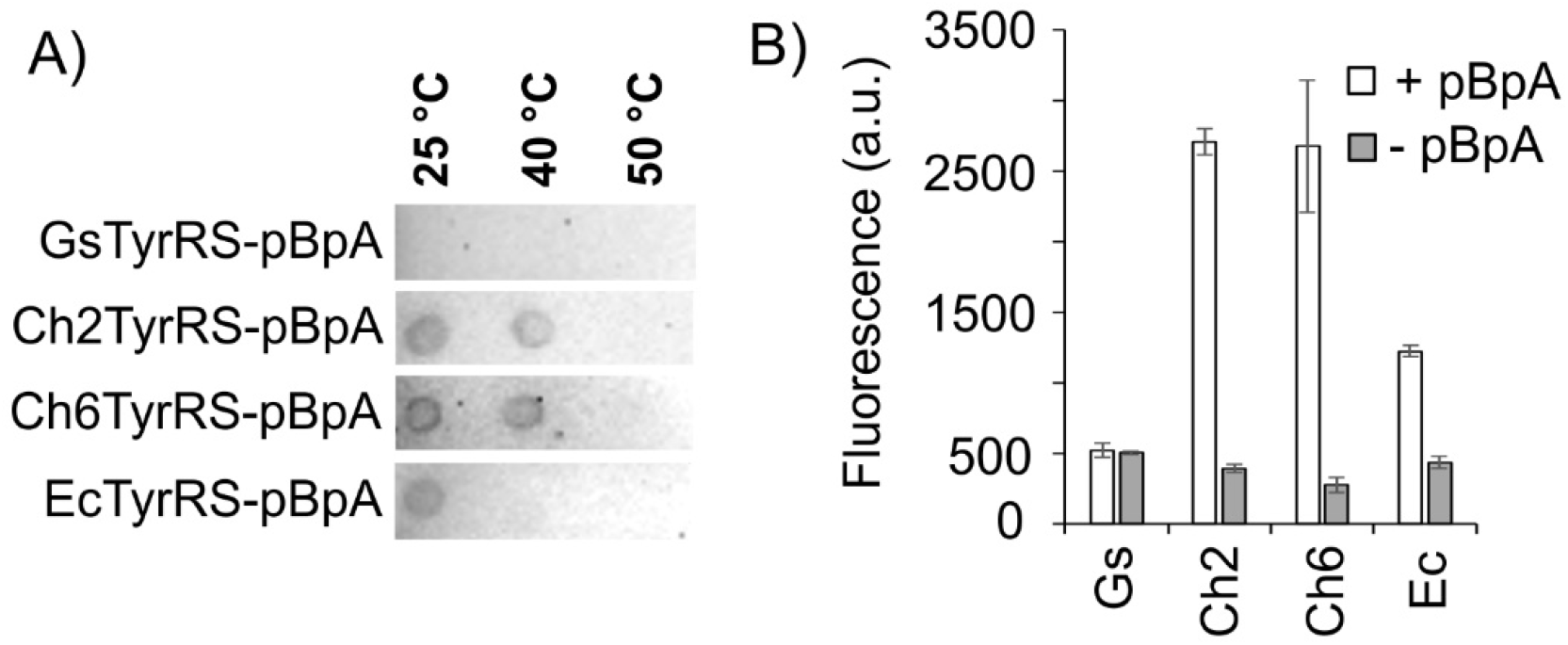
Chimeric TyrRS variants better tolerate pBpA-selective mutations. A) Thermal shift assay of the pBpA selective TyrRS mutants constructed from the two wild-type or chimeric scaffolds, in *E. coli* cell-free extract. Dot-blot was performed (using an anti-polyhistidine antibody) on the soluble fraction of *E. coli* cell-free extracts expressing the indicated TyrRS-pBpA variants (left), after incubation at the indicated temperature (top). B) Activity of the TyrRS-pBpA variants in the ATMY *E. coli* strain, measured using the expression of sfGFP-151TAG reporter in the presence of tRNA_CUA_^Tyr^ and in the presence and absence of 1 mM pBpA. Characteristic fluorescence of the full-length sfGFP-151-TAG reporter was measured in resuspended cells.

So far, we have evaluated the activity of these engineered pairs in our ATMY *E. coli* strain, where it is significantly easier to control parameters affecting the assay performance, such as the expression level of the aaRS/tRNA pairs. To demonstrate the utility of these new tools in eukaryotic cells, we tested the activity of the pBpA-selective TyrRS variants in HEK293T cells. EcTyrRS-pBpA, GsTyrRS-pBpA, Ch2TyrRS-pBpA, and Ch6TyrRS-pBpA were each cloned into a mammalian expression vector under a UbiC promoter, which also encodes 16 copies of the tRNA^Tyr^CUA expression cassette. The resulting plasmids were co-transfected into HEK293T cells with another plasmid encoding an EGFP-39-TAG reporter, and the full-length reporter expression was monitored in the presence or absence of 1 mM pBpA in the media (Figure 4A, Figure S4). Expression of a wild-type EGFP reporter (no in-frame TAG codon) was also included as a control. All four TyrRS-pBpA variants enabled successful incorporation of pBpA into the reporter. However, Ch2TyrRS-pBpA and Ch6TyrRS-pBpA demonstrated significantly higher activity (up to 36% of wild-type EGFP) relative to EcTyrRS-pBpA and GsTyrRS-pBpA (Figure 4A). The reporter protein expressed using Ch2TyrRS-pBpA was isolated using Ni-NTA affinity chromatography and characterized by SDS-PAGE (Figure S5) and mass-spectrometry analysis (Figure S6) to confirm successful pBpA incorporation. It is worth noting that the activity of GsTyrRS-pBpA in mammalian cells was found to be significantly higher than what is expected from its assessment in ATMY *E. coli*, where it was found to be nearly inactive. It is possible that the more sophisticated protein folding machinery of the mammalian cells is able to better process unstable engineered proteins like GsTyrRS-pBpA. The TyrRS variants are also expressed more strongly in mammalian cells than in *E. coli*, which may also contribute to the observed difference. Nonetheless, these results demonstrate that the chimeric TyrRS variants provide improved platforms for the GCE technology in eukaryotes. To further highlight the generality of this approach, we constructed Ch2TyrRS-Poly by introducing previously reported active site mutations for generating EcTyrRS-Poly (Figure 1A), an engineered mutant that charges several different ncAAs including O-methyltyrosine (OMeY). When tested in HEK293T cells, both EcTyrRS-Poly and Ch2TyrRS-Poly exhibited comparable activity for EGFP-39-TAG reporter expression (Figure 4B, Figure S4), confirming that the chimeric TyrRS is able to recapitulate the activity of well-behaved bacterial engineered EcTyrRS mutants, while providing a better scaffold for those with suboptimal stability.

**Figure 4.**
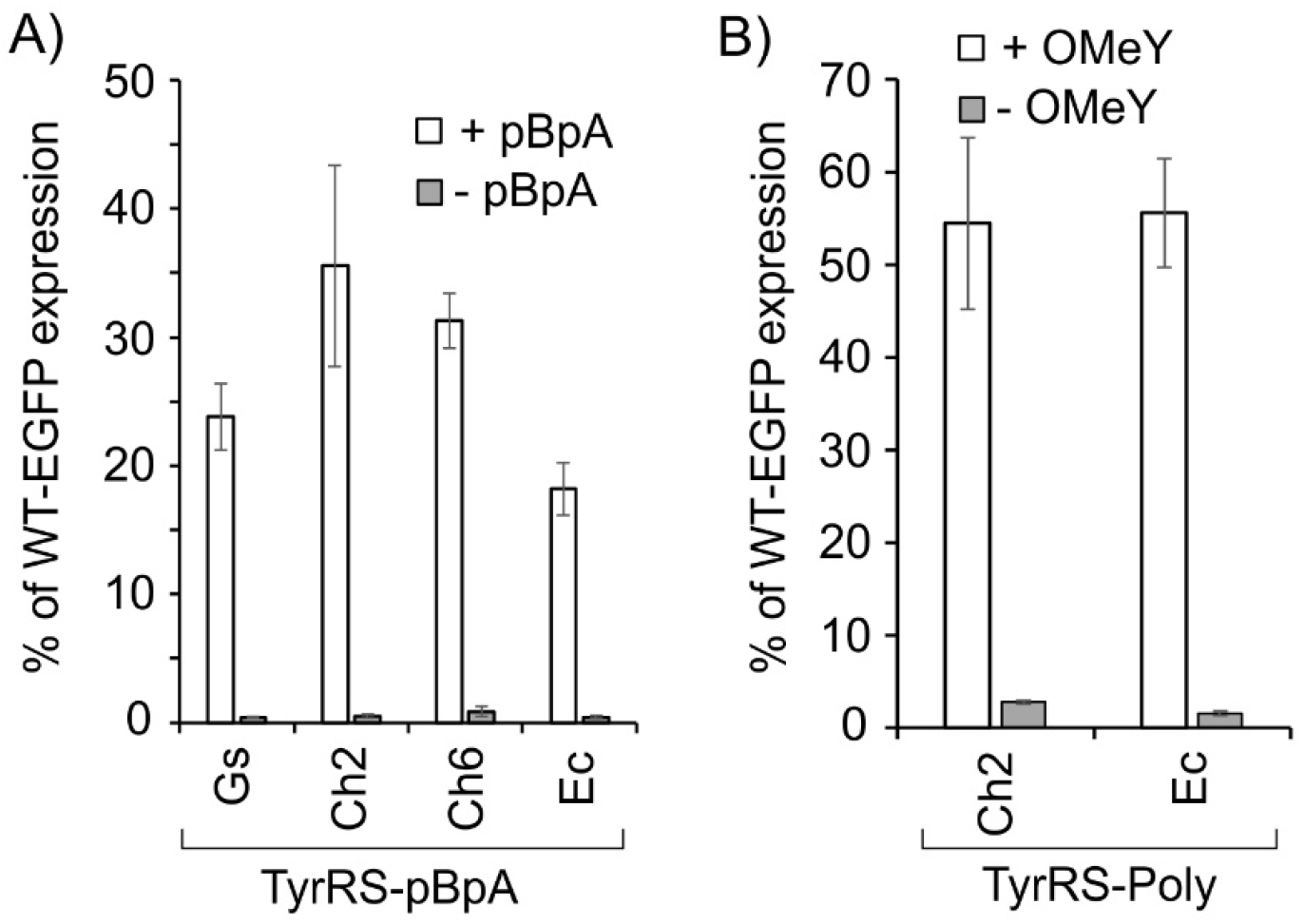
Chimeric TyrRS derived mutants exhibit improved activity for ncAA incorporation in mammalian cells. A) TyrRS-pBpA variants were co-expressed in HEK293T cells with tRNA_CUA_^Tyr^ and EGFP-39-TAG, in the presence and absence of pBpA, and the expression of the full-length reporter was monitored by its characteristic fluorescence in cell-free extract. The expression level was reported as the % of an identical experiment where wild-type EGFP reporter (no TAG) was used as the reporter. B) The ncAA-incorporation efficiency of Ch2TyrRS-Poly is comparable to the highly active EcTyrRS-Poly. Activity was measured as described in section (A) in the presence and absence of 1 mM OMeY. See Figure S4 for all associated fluorescence images.

Work in the last two decades have provided a deep insight into how the biophysical properties of a protein influence its evolution.^*18–21*^ It is now clear that the function-altering mutations acquired during experimental or natural evolution can often negatively impact the structural stability of a protein. Our work highlights its impact on the GCE technology, which relies on engineered aaRSs that selectively charge ncAAs of interest. We show how the structural robustness of MjTyrRS has contributed to its remarkable success as a powerful GCE platform. In contrast, the lower stability of EcTyrRS compromises the extent to which its active site can be altered. It is important to note that the same limitation likely affects the engineerability of several other aaRS/tRNA pairs, derived from mesophilic organisms (e.g., *E. coli* or yeast),^*3, 6, 33–35*^ which have been adapted for ncAA incorporation. Indeed, the success of engineering these platforms for ncAA incorporation has been limited. We describe a strategy to overcome this challenge by taking advantage of more thermostable aaRS homologs derived from thermophilic organisms. We further show that chimeras generated from thermophilic and mesophilic aaRS homologs may be even better suited for this purpose. Analogous strategies have been used to create optimal starting points for the directed evolution of enzymes such as cytochrome P450.^*31*^ It is possible that instead of simple aaRS chimeras like the ones reported here, more sophisticated counterparts with even better properties can be created by constructing and selecting a DNA-shuffling library of EcTyrRS and GsTyrRS. Additionally, we find that relative performance of the same aaRS in different expression systems can be different. For example, the GsTyrRS-pBpA mutant, which is largely insoluble and inactive in *E. coli*, demonstrated robust activity in mammalian cells. This might stem from differences in protein folding machinery in different host cells, as well as other factors such as variable expression level of the aaRS, speed of translation, codon usage, etc.

In summary, here we establish the structural robustness of an aaRS as an important factor that significantly impacts its engineerability for GCE. We also provide a roadmap for creating more engineerable bacterial aaRS variants by hybridizing homologs from mesophilic and thermophilic bacteria. Mutants generated from such chimeric TyrRSs show robust activity in both ATMY *E. coli* strain as well as in mammalian cells, suggesting that these are more attractive scaffolds for extensive engineering. Directed evolution of these using our facile ATMY *E. coli* based selection system should provide access to new enabling ncAAs. Finally, improved pBpA-incorporation activity of ChTyrRS-pBpA will further facilitate the application of this important photo-crosslinker ncAA for uncovering new biomolecular interactions in eukaryotic cells.

## Notes

The authors declare no competing financial interest.

## ACKNOWLEDGMENTS

This work was partially supported by NIH grants R01GM126220 and R01GM124319 to A.C.

---

**Figure S1:**
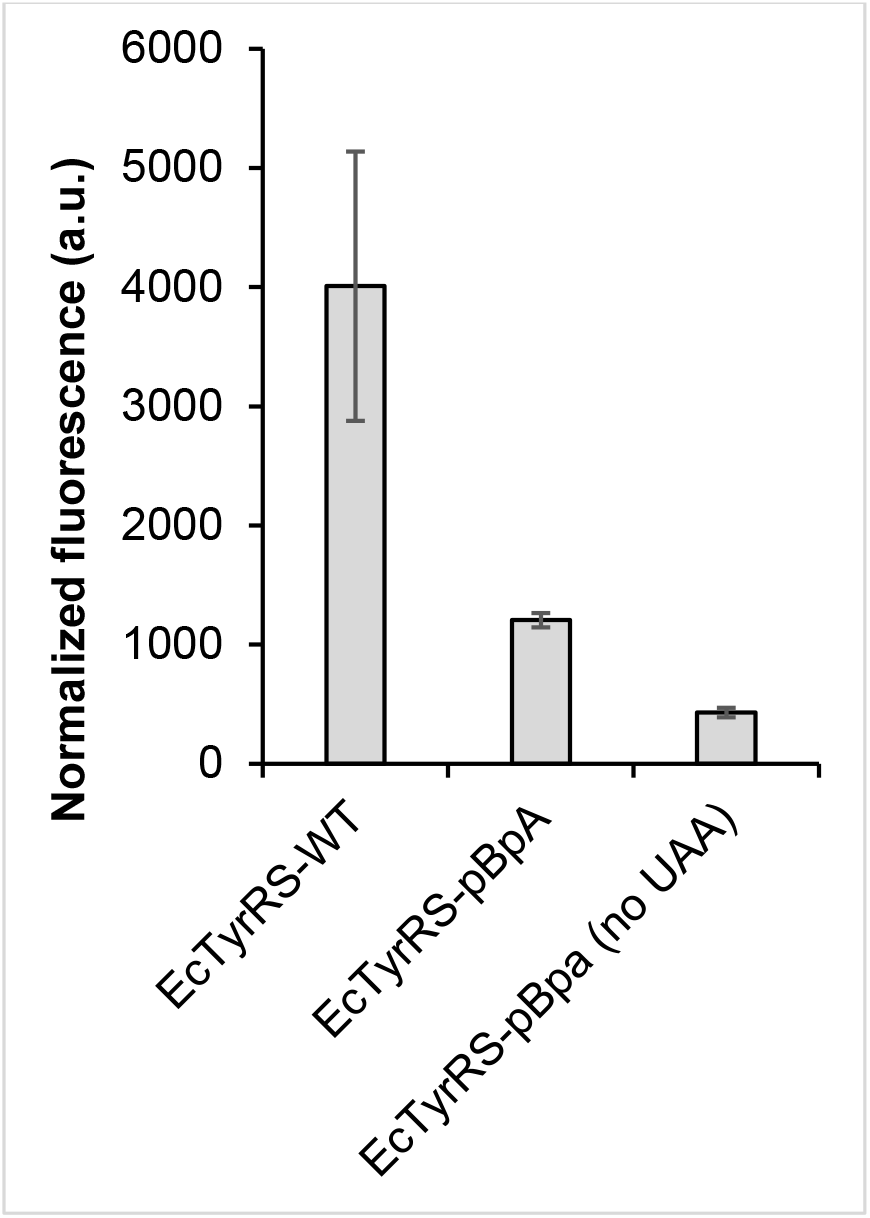
The activity of EcTyrRS-pBpA is significantly weaker than EcTyrRS-WT. Activity was evaluated using the sfGFP-151-TAG reporter, expressed in ATMY *E. coli*, by measuring the characteristic fluorescence of the full-length reporter in resuspended cells. For EcTyrRS-pBpA, the expression was measured in the presence or absence of 1 mM pBpA.

**Figure S2:**
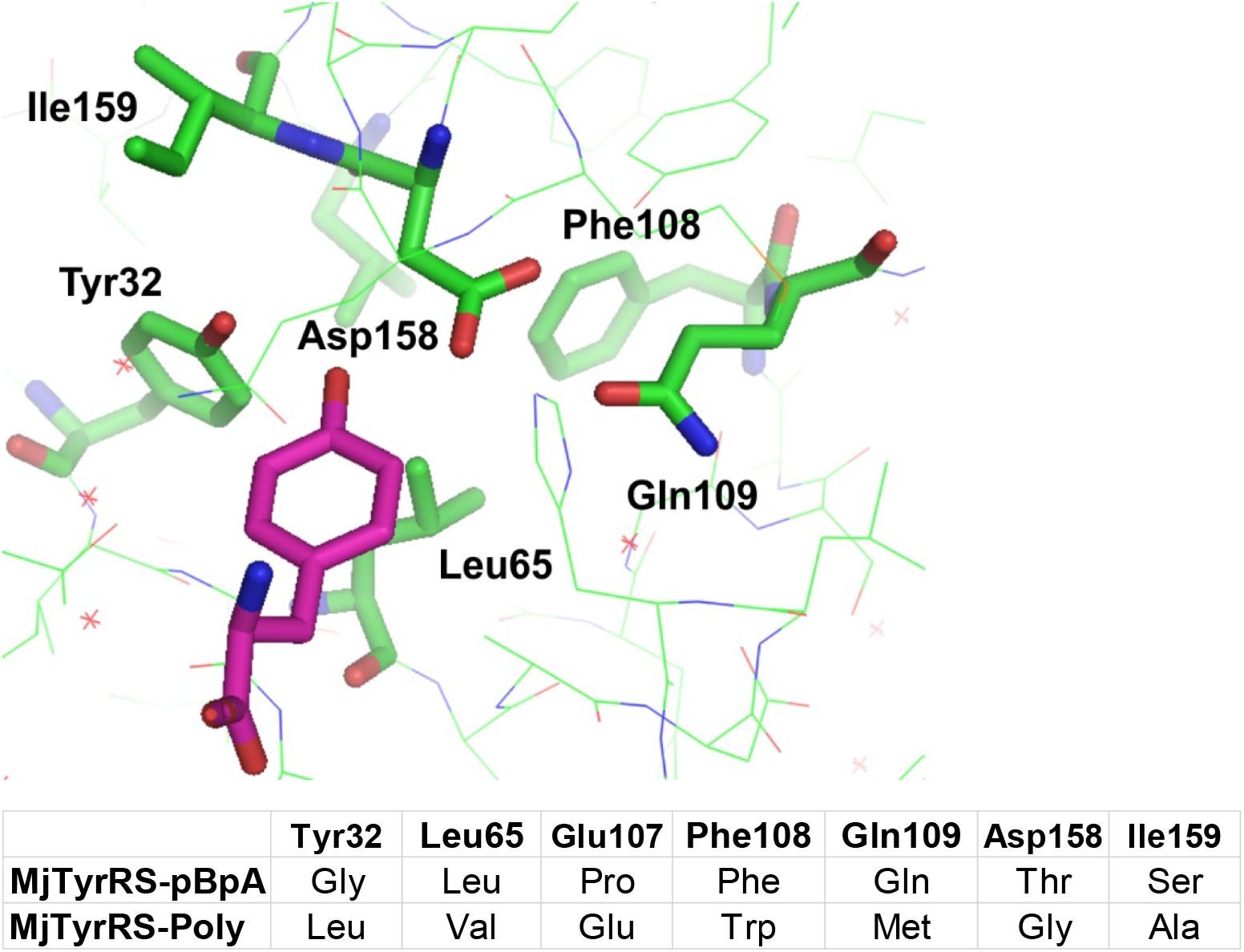
MjTyrRS mutants used in this study. The active site structure of MjTyrRS is also shown highlighting the key residues mutated to generate ncAA-selective variants.

**Figure S3.**
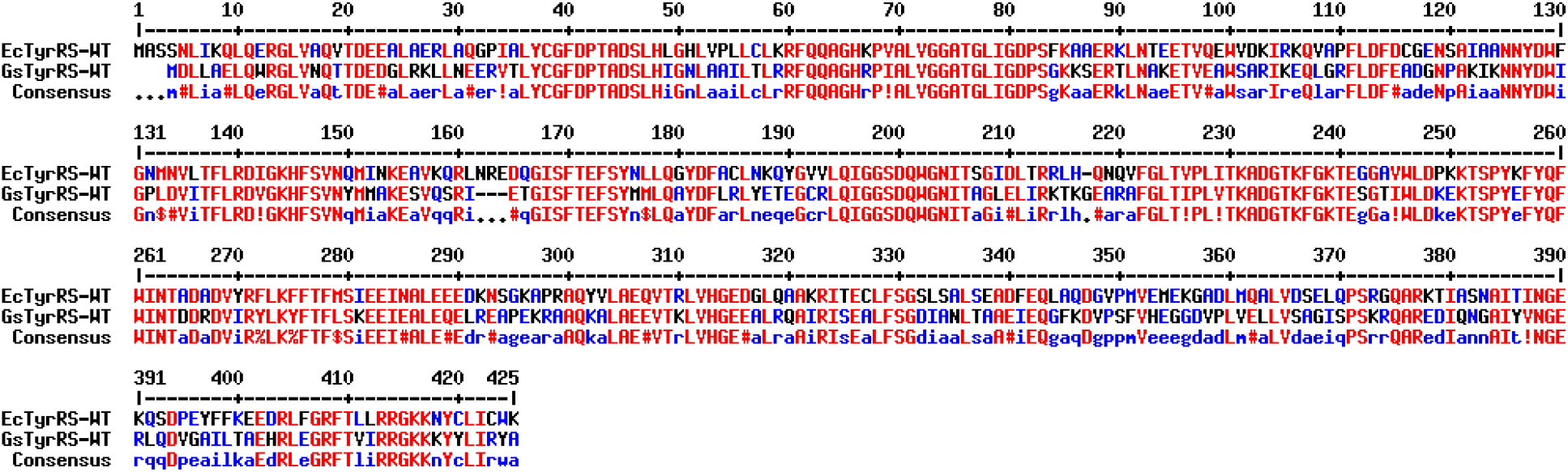
Sequence alignment of EcTyrRS-WT and GsTyrRS-WT.

**Figure S4.**
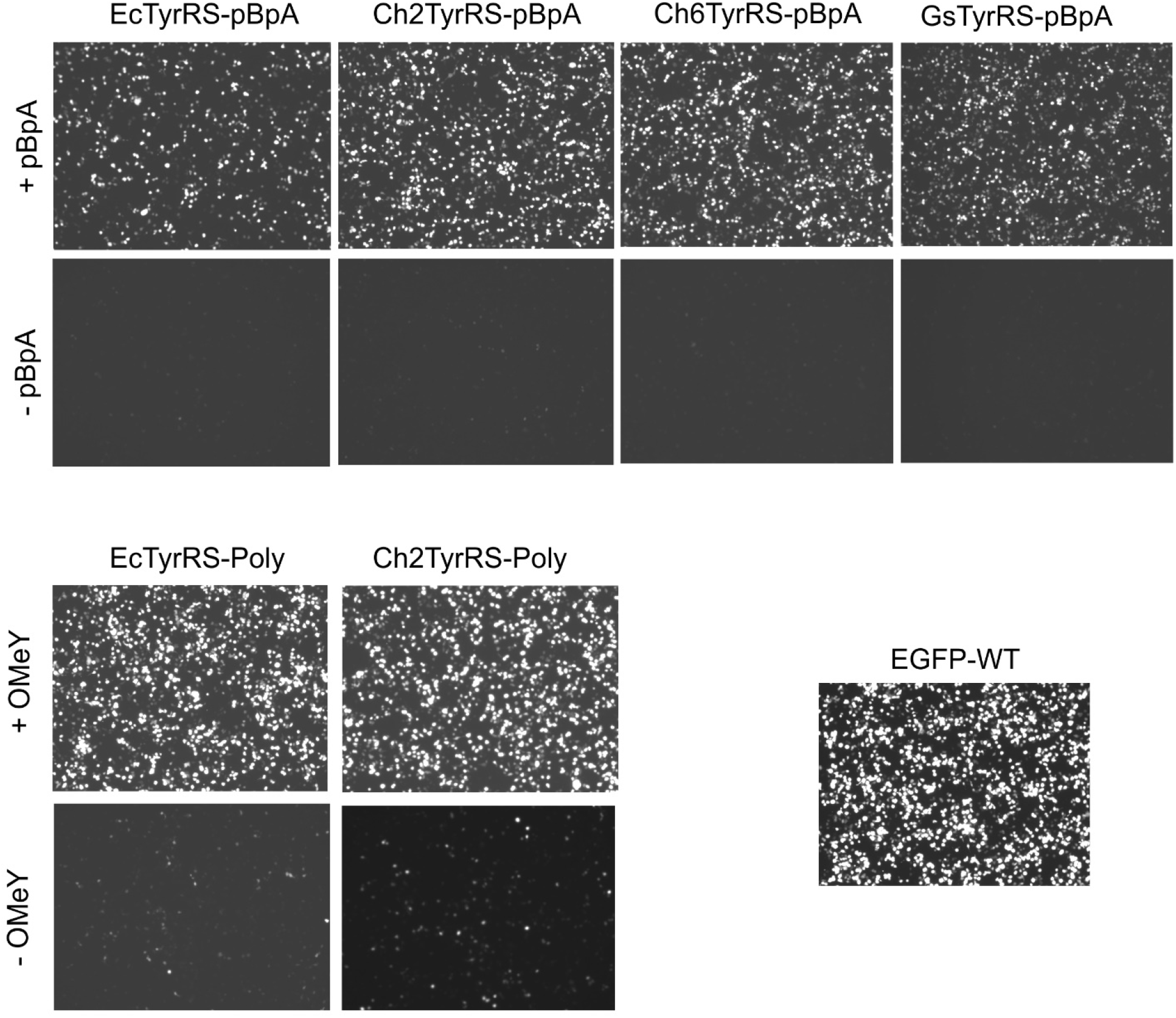
Fluorescence images of mammalian cells expressing EGFP-39-TAG reporter using various TyrRS/tRNA pairs (shown above), in the presence or absence of relevant ncAAs (left). These images correspond to the fluorescence values presented in Figure 4.

**Figure S5.**
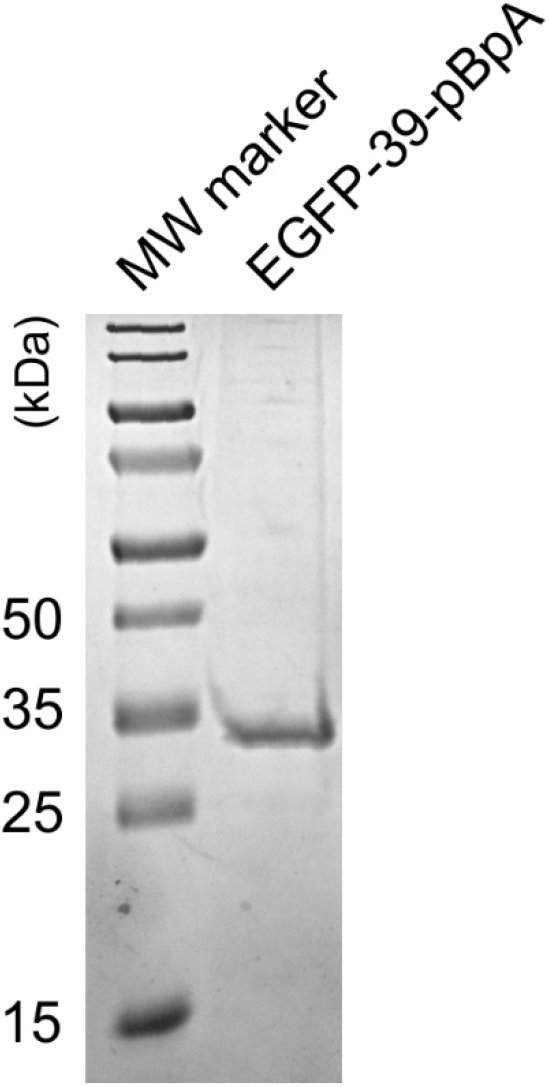
SDS-PAGE analysis of the EGFP-39-pBpA reporter expressed in HEK293T cells using Ch2TyrRS-pBpA.

**Figure S6.**
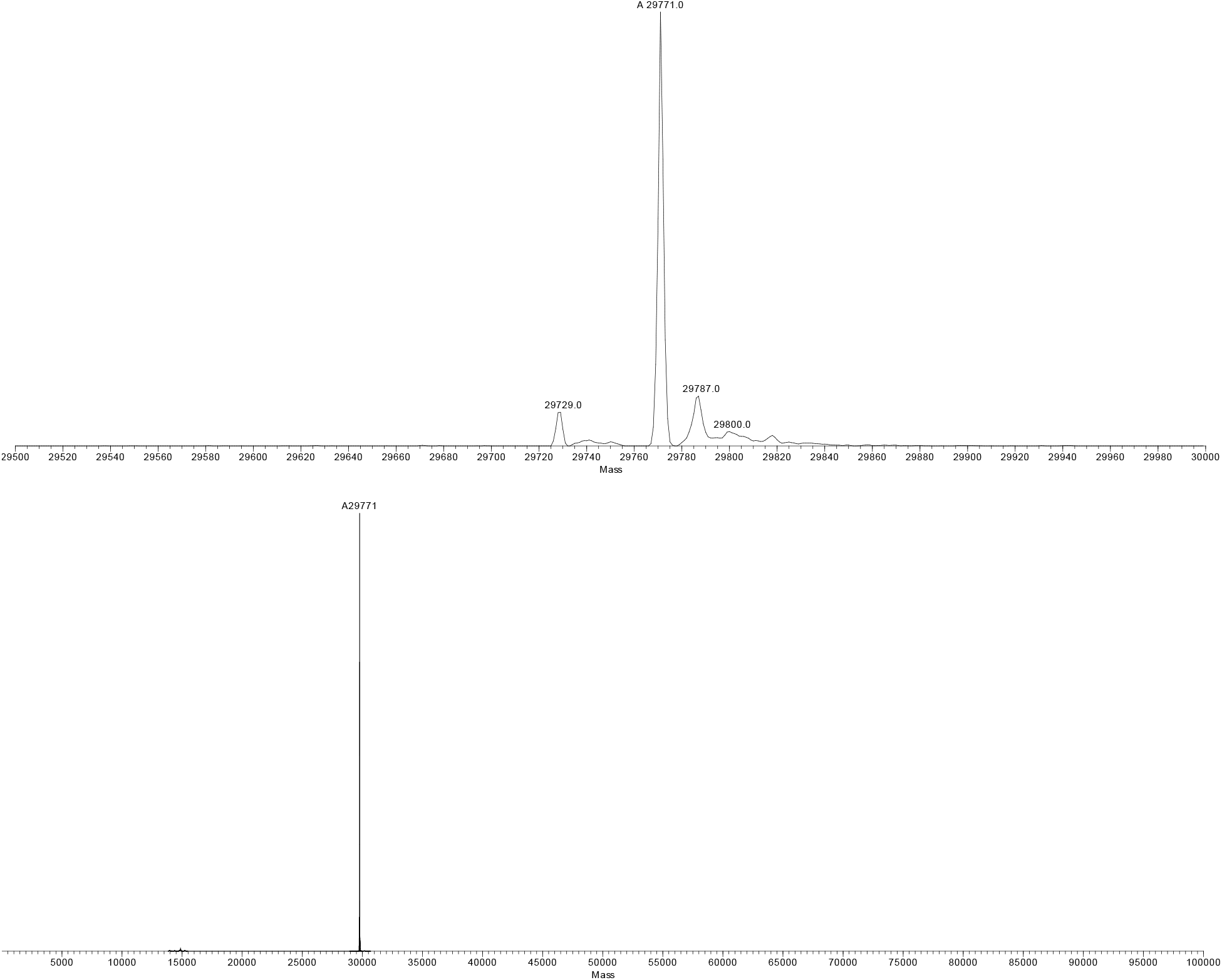
Deconvoluted ESI-MS analysis of EGFP-39-pBpA reporter expressed in HEK293T cells using Ch2TyrRS-pBpA. Two different magnifications of the same spectra are shown. The major species (29771 Da) corresponds to the expected mass; 29729 Da peak likely represents the same species lacking N-terminal acetylation (− 42 Da), while 29787 Da peak likely arises from oxidation (+16 Da).

