## Supplementary material for "Structural robustness affects the engineerability of aminoacyl-tRNA synthetases for genetic code expansion": Materials and methods

### General materials:

All cloning and plasmid propagation were done in DH10B *E. coli* cells. Restriction enzymes, Phusion HS II High-Fidelity DNA polymerase, and IPTG were obtained from Fisher. T4 DNA ligase was obtained from Enzymatics. DNA extraction and PCR clean up were conducted with Macherey-Nagel Binding Buffer NTI and Epoch mini spin columns from Thermo Fisher Scientific. Media components were obtained from Fisher Scientific. The following antibiotic stock concentrations were used: ampicillin 100 μg/mL, kanamycin 50 μg/mL, spectinomycin 100 μg/mL, chloramphenicol 35 μg/mL for LB Agar plates and cultures. A Cole Parmer Ultrasonic Processor was used for making *E. coli* lysate by sonication. Protein purification was conducted with a HisPur Ni-NTA resin from ThermoScientific. Dot blots were done with GE Healthcare life sciences nitrocellulose blotting membrane (0.45 um). Western blots were conducted with a PVDF membrane, antibodies, and SuperSignal West Dura Extended Duration Substrate for western blot from Thermo Fisher Scientific.

### Bacterial and virus strains

| Strain | Source | Catalog Number |
|---|---|---|
| ATMY6 | Chatterjee lab | N/A |
| *E. coli* DH10B | Thermo Fisher Scientific | 18297010 |

### Chemicals, peptides, and recombinant proteins

| Reagent | Source | Catalog Number |
|---|---|---|
| para-benzoyl-l-phenylalanine | Chem-Impex International | 05110 |
| O-methyl tyrosine | Fisher Scientific | AAH6309606 |

### Experimental models: cell lines

| Strain | Source | Catalog Number |
|---|---|---|
| HEK293T | ATCC | CRL-1573 |

**Construction of plasmids to express aaRS for ncAA incorporation into sfGFP and EGFP:** The *G. stearothermophilus* tyrosyl-aaRS was PCR amplified from a gBlock purchased from IDT, digested with NdeI/NcoI, and inserted into the pBK vector backbone. The mutant *G. stearothermophilus* tyrosyl-aaRS mutants were then generated via standard site-directed mutagenesis of the appropriate active site residues. The pBK *E. coli* tyrosyl-aaRS wild type and mutants were previously reported.[1,2]

The chimeric H2 and H6 aaRS' were constructed by PCR amplification of the *E. coli* tyrosyl-aaRS N-terminus and the *G. stearothermophilus* C-terminus. The inserts were then made through overlap amplification, digested with NdeI/NcoI, and inserted into the pBK vector backbone. The mutant chimeric aaRS were then generated via standard site-directed mutagenesis.[3]

For expression of the aaRS in mammalian cells, terminal primers were used to PCR amplify the aaRS' from their respective pBK plasmids. This was followed by digestion of the PCR products with NheI/XhoI into the pB1U vector backbone.

**Construction of plasmids to express aaRS for CETSA:** The *E. coli*, *G. stearothermophilus*, and chimeric aaRS' were PCR amplified from their respective pBK constructs, digested with NdeI/HindIII, and inserted into the pET22b vector backbone. The N-terminal primer appended a 10X-Histidine tag to each aaRS for future imaging.

**sfGFP* fluorescence analysis and expression:** For *E. coli* expression, the pBK aaRS and pEvol T5 EcY-TAG sfGFP151* reporter plasmids were co-transformed into ATMY6 cells.[1] A 5 mL overnight culture was inoculated with a single colony and the appropriate antibiotics. The overnight starter culture was then used to inoculate a 20 mL LB Media culture supplemented with antibiotics. Cultures were grown to an OD600 of 0.6 then induced with a final concentration of 1 mM IPTG, the appropriate ncAA (1 mM), and incubated for 16 hours at 30 $^{o}$C with shaking (250 rpm). The cultures were then spun down, the LB media was removed, and the cells were resuspended in 1X PBS. Fluorescence readings were collected in a 96-well plate using a SpectraMAX M5 (Molecular Devices) (ex= 488 nm and em=534 nm). Mean of two independent experiments were reported, and error bars represent standard deviation.

**EGFP* fluorescence analysis, expression and purification:** For fluorescence analysis, HEK 293T cells were seeded at a density of 600,000 cells per well for a 12-well plate the day before transfection. A total amount of 1.5 μg DNA (0.75 μg of each plasmid for two-plasmids) + 3.5 μL PEI + 17.5 μL DMEM was used for transfection of each well. Fluorescence images and EGFP expression analysis were performed 48 hours post transfection following previously mentioned protocols.[4] Fluorescence readings were collected in a 96-well plate using a SpectraMAX M5 (Molecular Devices) (ex= 488 nm and em=510 nm). Mean of four independent experiments were reported, and error bars represent standard deviation.

For EGFP protein purification incorporating one ncAA, HEK293T cells were seeded in 100 mm cell culture dishes (5 million per dish) 24 hours prior to transfection.

**CETSA assay:** For aaRS expression, TOP10 *E. coli* cells were transformed with a single pET22b-N-terminal-10X-Histidine tagged-aaRS plasmid. Overnight cultures were inoculated with a single colony, then used to inoculate 20 mL LB Media cultures with the appropriate antibiotics, grown to an OD600 of 0.6, and induced with IPTG (final concentration of 1 mM) for 15 minutes at 30 $^{o}$C with shaking. The cultures were then spun down, the LB Media was removed, and the cell pellets were resuspended in 500 μL of sonication buffer (100 mM NaCl, 25 mM Tris HCl, pH 8.0).

For lysate preparation, the cell pellets were treated to three freeze thaw cycles followed by three cycles of sonication (75% power, 20 pulses), spun down, and the supernatant was collected. Each supernatant was divided into 50 μL aliquots and heated at varying temperatures for 5 minutes on a Perkin Elmer Cetus DNA Thermal Cycler 480 and spun down at maximum speed for 10 minutes. Then 3 μL of the supernatant was inoculated on a nitrocellulose membrane and treated to western blot analysis following previously described protocols.[4] Antibodies used for imaging include: mouse anti-Histidine 6X tag antibody (1:1000 dilution), chicken anti-mouse IgG secondary antibody-HRP conjugate (1:5000 dilution).

**Solubility Western:** DH10B cells were transformed with a pET22b-N-term-10X-His-aaRS plasmid and an overnight 5 mL culture was inoculated with a starter colony. A 20 mL culture was then inoculated and grown to an OD600 of 0.6, allowed to grow for 4 hours at 30 $^{o}$C, spun down, and lysed by sonication following the same protocol as the aforementioned CETSA assay. This was resolved using 12% SDS-PAGE gel and worked up for a western following previously described protocols.[4] The antibodies used for this protocol were the same as those for the CETSA assay.

**Primers and other DNA sequences:**

EcYRS NdeI-F
TTTGAGGAATCCCATATGGCAAGCAGTAACTTGATTAAACAATTGCAAGAG

EcYRS NcoI-R
AATTCCATGGTTATTTCCAGCAAATCAGACACTAATTC

GsYRS NdeI-F
ATTATTATGAATCCCATATGATGGACCTGCTGGCGGAACTGCAATG

pBK MCS JI sq-R
GAGATCATGTAGGCCTGATAAGCGTAGC

H2 EcYRS-iR
GCGATCGGACGGTGACCCGCCTGCTGGAAGCGTTTCAGGCATAACAATG

H2 GsYRS-iF
CCTGAAACGCTTCCAGCAGGCGGGTCACCGTCCGATCGCGCTGGTTG

H6 EcYRS-iR
CCGCTTTCGGTTTTGCCAAATTTGGTGCCATCTGCTTTAGTGATCAGCGGAAC
G

H6 GsYRS-iF
CACTAAAGCAGATGGCACCAAATTTGGCAAAACCGAAAGCGGTACCATTTG

EcYRS NheI-F
TTTGAGGAATCCGCTAGCGCAAGCAGTAACTTGATTAAACAATTGCAAGAG

EcYRS XhoI-R
AATTCTCGAGTTATTTCCAGCAAATCAGACACTAATTC

GsYRS NheI-F
AATAATGCTAGCATGGACCTGCTGGCG

GsYRS XhoI-R
AATTCTCGAGTTACGCATAACGAATCAGATAGTATTTC

EcYRS-nterm10XHis-NdeI-F
GAAATTACATATGCATCATCACCATCACCATCATCATCATCACGCAAGCAGT
AACTTGATTAAACAATTGCAAGAG

EcYRS-HindIII-R
AATTAAGCTTTTATTTCCAGCAAATCAGACACTAATTC

GsYRS-nterm10XHis-NdeI-F
GAAATTACATATGCATCATCACCATCACCATCATCATCATCACGACCTGCTGG
CGGAACTGCAATGG

GsYRS-HindIII-R
AATTAAGCTTTTACGCATAACGAATCAGATAGTATTTC

MjYRS-nterm10XHis-NdeI-F
GAAATTACATATGCATCATCACCATCACCATCATCATCATCACgacgaatttgaaatgat
aaagagaaacacatctg

MjYRS-HindIII-R
AATTAAGCTTTTATAATCTCTTTCTAATTGGCTCTAAAATC

GeobacYRS-Y34G-R
GCTATCCGCGGTCGGGTCGAAACCGCAACCCAGGGTCACACGTTCCTCGTTC
AGC

GeobacYRS-D176G-R
CAGCCTTCGGTTTCGTACAGACGCAGGAAACCATACGCTTGCAGCATCATGT
AGCTAAAC

GeobacYRS-GGFL-L180A-R
CAGACGGCAGCCTTCGGTTTCGTAGGCACGCAGGAAACCATACGCTTG

GeobacYRS-D176G-F

GTTTAGCTACATGATGCTGCAAGCGTATGGTTTCCTGCGTCTGTACGAAACCG
AAGGCTG

GeobacYRS-GGFL-L180A-F
CAAGCGTATGGTTTCCTGCGTGCCTACGAAACCGAAGGCTGCCGTCTG

gBlock sequence of *G. stearothermophilus* tyrosyl aminoacyl-tRNA synthetase
ATGGCGAGCAGCGACCTGCTGGCGGAACTGCAATGGCGTGGCCTGGTTAATC
AGACCACCGACGAAGATGGCCTGCGTAAACTGCTGAACGAGGAACGTGTGA
CCCTGTATTGCGGTTTCGACCCGACCGCGGATAGCCTGCACATCGGCAACCT
GGCGGCGATTCTGACCCTGCGTCGTTTTCAGCAAGCGGGTCACCGTCCGATC
GCGCTGGTTGGTGGTGCGACCGGTCTGATTGGCGACCCGAGCGGCAAGAAAA
GCGAGCGTACCCTGAACGCGAAGGAAACCGTTGAAGCGTGGAGCGCGCGTA
TCAAAGAACAGCTGGGTCGTTTCCTGGACTTTGAGGCGGATGGCAACCCGGC
GAAGATTAAAAACAACTATGACTGGATCGGTCCGCTGGATGTGATTACCTTC
CTGCGTGATGTGGGCAAGCACTTTAGCGTTAACTACATGATGGCGAAAGAGA
GCGTTCAGAGCCGTATCGAAACCGGTATTAGCTTCACCGAGTTTAGCTACATG
ATGCTGCAAGCGTATGACTTCCTGCGTCTGTACGAAACCGAAGGCTGCCGTCT
GCAGATCGGTGGCAGCGATCAATGGGGTAACATCACCGCGGGCCTGGAACTG
ATTCGTAAGACCAAAGGTGAAGCGCGTGCGTTTGGCCTGACCATCCCGCTGG
TGACCAAGGCGGACGGTACCAAGTTTGGCAAAACCGAAAGCGGTACCATTTG
GCTGGATAAGGAGAAAACCAGCCCGTACGAATTCTATCAGTTTTGGATCAAC
ACCGACGATCGTGACGTTATTCGTTACCTGAAGTATTTCACCTTTCTGAGCAA
AGAGGAAATCGAAGCGCTGGAGCAGGAACTGCGTGAGGCGCCGGAAAAGCG
TGCGGCGCAAAAAGCGCTGGCGGAGGAAGTGACCAAACTGGTTCACGGTGA
GGAAGCGCTGCGTCAGGCGATCCGTATTAGCGAAGCGCTGTTTAGCGGTGAT
ATCGCGAACCTGACCGCGGCGGAGATTGAACAAGGCTTCAAGGACGTGCCG
AGCTTTGTTCACGAAGGTGGCGATGTGCCGCTGGTTGAGCTGCTGGTTAGCGC
GGGTATCAGCCCGAGCAAACGTCAGGCGCGTGAAGACATCCAAAACGGTGC
GATTTACGTGAACGGCGAGCGTCTGCAAGATGTTGGCGCGATTCTGACCGCG
GAACACCGTCTGGAAGGTCGTTTTACCGTTATCCGTCGTGGCAAGAAGAAAT
ACTATCTGATTCGTTATGCGTAA

**Plasmids:** The text of the final plasmid maps and sequences are provided below with the following color coding: aaRS highlighted red, antibiotic selectable marker highlighted blue, tRNA highlighted purple, lacI highlighted green, and the origin of replication highlighted orange. The images are not color coded.

pBK MCS GsYRS wt

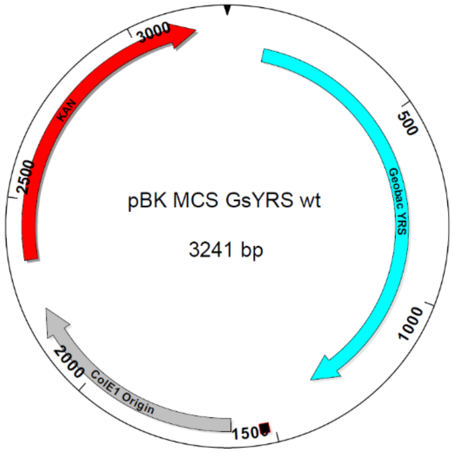


cttttgctgagttgaaggatccgcggccgctcgggttgtcagcctgtcccgcttataagatcatacgccgttatacgttgtttacgctttgagga
atcccatatgATGGACCTGCTGGCGGAACTGCAATGGCGTGGCCTGGTTAATCAGACCACC
GACGAAGATGGCCTGCGTAAACTGCTGAACGAGGAACGTGTGACCCTGTATTGCGG
TTTCGACCCGACCGCGGATAGCCTGCACATCGGCAACCTGGCGGCGATTCTGACCCT
GCGTCGTTTTCAGCAAGCGGGTCACCGTCCGATCGCGCTGGTTGGTGGTGCGACCGG
TCTGATTGGCGACCCGAGCGGCAAGAAAAGCGAGCGTACCCTGAACGCGAAGGAA
ACCGTTGAAGCGTGGAGCGCGCGTATCAAAGAACAGCTGGGTCGTTTCCTGGACTTT
GAGGCGGATGGCAACCCGGCGAAGATTAAAAACAACTATGACTGGATCGGTCCGCT
GGATGTGATTACCTTCCTGCGTGATGTGGGCAAGCACTTTAGCGTTAACTACATGAT
GGCGAAAGAGAGCGTTCAGAGCCGTATCGAAACCGGTATTAGCTTCACCGAGTTTA
GCTACATGATGCTGCAAGCGTATGACTTCCTGCGTCTGTACGAAACCGAAGGCTGCC
GTCTGCAGATCGGTGGCAGCGATCAATGGGGTAACATCACCGCGGGCCTGGAACTG
ATTCGTAAGACCAAAGGTGAAGCGCGTGCGTTTGGCCTGACCATCCCGCTGGTGACC
AAGGCGGACGGTACCAAGTTTGGCAAAACCGAAAGCGGTACCATTTGGCTGGATAA
GGAGAAAACCAGCCCGTACGAATTCTATCAGTTTTGGATCAACACCGACGATCGTG
ACGTTATTCGTTACCTGAAGTATTTCACCTTTCTGAGCAAAGAGGAAATCGAAGCGC
TGGAGCAGGAACTGCGTGAGGCGCCGGAAAAGCGTGCGGCGCAAAAAGCGCTGGC
GGAGGAAGTGACCAAACTGGTTCACGGTGAGGAAGCGCTGCGTCAGGCGATCCGTA
TTAGCGAAGCGCTGTTTAGCGGTGATATCGCGAACCTGACCGCGGCGGAGATTGAA
CAAGGCTTCAAGGACGTGCCGAGCTTTGTTCACGAAGGTGGCGATGTGCCGCTGGTT
GAGCTGCTGGTTAGCGCGGGTATCAGCCCGAGCAAACGTCAGGCGCGTGAAGACAT
CCAAAACGGTGCGATTTACGTGAACGGCGAGCGTCTGCAAGATGTTGGCGCGATTCT
GACCGCGGAACACCGTCTGGAAGGTCGTTTTACCGTTATCCGTCGTGGCAAGAAGA
AATACTATCTGATTCGTTATGCGTAAggtaccatggctgcagtttcaaacgctaaattgcctgatgcgctacgcttat
caggcctacatgatctctgcaatatattgagtttgcgtgcttttgtaggccggataaggcgttcacgccgcatccggcaagaaacagcaaaca
atccaaaacgccgcgttcagcggcgtttttctgcttttcttcgcgaattaattccgcttcgcacatgtgagcaaaaggccagcaaaaggccag
gaaccgtaaaaaggccgcgttgctggcgtttttccataggctccgcccccctgacgagcatcacaaaaatcgacgctcaagtcagaggtgg
cgaaacccgacaggactataaagataccaggcgtttccccctggaagctccctcgtgcgctctcctgttccgaccctgccgcttaccggata
cctgtccgcctttctcccttcgggaagcgtggcgctttctcatagctcacgctgtaggtatctcagttcggtgtaggtcgttcgctccaagctgg
gctgtgtgcacgaacccccgttcagcccgaccgctgcgccttatccggtaactatcgtcttgagtccaacccggtaagacacgacttatcg

ccactggcagcagccactggtaacaggattagcagagcgaggtatgtaggcggtgctacagagttcttgaagtggtggcctaactacggct
acactagaaggacagtatttggtatctgcgctctgctgaagccagttaccttcggaaaaagagttggtagctcttgatccggcaaacaaacca
ccgctggtagcggtggtttttttgtttgcaagcagcagattacgcgcagaaaaaaaggatctcaagaagatcctttgatcttttctacggggtct
gacgctcagtggaacgaaaactcacgttaagggattttggtcatgaacaataaaactgtctgcttacataaacagtaatacaaggggtgttatg
agccatattcaacgggaaacgtcttgctcgaggccgcgattaaattccaacatggatgctgatttatatgggtataaatgggctcgcgataatg
tcgggcaatcaggtgcgacaatctatcgattgtatgggaagcccgatgcgccagagttgtttctgaaacatggcaaaggtagcgttgccaat
gatgttacagatgagatggtcagactaaactggctgacggaatttatgcctcttccgaccatcaagcattttatccgtactcctgatgatgcatg
gttactcaccactgcgatccccgggaaaacagcattccaggtattagaagaatatcctgattcaggtgaaaatattgttgatgcgctggcagt
gttcctgcgccggttgcattcgattcctgtttgtaattgtccttttaacagcgatcgcgtatttcgtctcgctcaggcgcaatcacgaatgaataac
ggtttggttgatgcgagtgattttgatgacgagcgtaatggctggcctgttgaacaagtctggaaagaaatgcataagcttttgccattctcacc
ggattcagtcgtcactcatggtgatttctcacttgataaccttatttttgacgaggggaaattaataggttgtattgatgttggacgagtcggaatc
gcagaccgataccaggatcttgccatcctatggaactgcctcggtgagttttctccttcattacagaaacggctttttcaaaaatatggtattgat
aatcctgatatgaataaattgcagtttcatttgatgctcgatgagtttttctaatcagaattggttaattggttgtaacactggcagagcattacgct
gacttgacgggacggcggctttgttgaataaatcgaa

pBK MCS Gs-BPA-RS
Same as above but with the following active site mutations:
Y34G, D176G, L180A

pBK MCS Chimera H2-YRS
Same as above but with the following aaRS sequence:
atgGCAAGCAGTAACTTGATTAAACAATTGCAAGAGCGGGGGCTGGTAGCCCAGGTG
ACGGACGAGGAAGCGTTAGCAGAGCGACTGGCGCAAGGCCCGATCGCGCTCTATTG
CGGCTTCGATCCTACCGCTGACAGCTTGCATTTGGGGCATCTTGTTCCATTGTTATGC
CTGAAACGCTTCCAGCAGGCGGGTCACCGTCCGATCGCGCTGGTTGGTGGTGCGACC
GGTCTGATTGGCGACCCGAGCGGCAAGAAAAGCGAGCGTACCCTGAACGCGAAGG
AAACCGTTGAAGCGTGGAGCGCGCGTATCAAAGAACAGCTGGGTCGTTTCCTGGAC
TTTGAGGCGGATGGCAACCCGGCGAAGATTAAAAACAACTATGACTGGATCGGTCC
GCTGGATGTGATTACCTTCCTGCGTGATGTGGGCAAGCACTTTAGCGTTAACTACAT
GATGGCGAAAGAGAGCGTTCAGAGCCGTATCGAAACCGGTATTAGCTTCACCGAGT
TTAGCTACATGATGCTGCAAGCGTATGATTTCCTGCGTCTGTACGAAACCGAAGGCT
GCCGTCTGCAGATCGGTGGCAGCGATCAATGGGGTAACATCACCGCGGGCCTGGAA
CTGATTCGTAAGACCAAAGGTGAAGCGCGTGCGTTTGGCCTGACCATCCCGCTGGTG
ACCAAGGCGGACGGTACCAAGTTTGGCAAAACCGAAAGCGGTACCATTTGGCTGGA
TAAGGAGAAAACCAGCCCGTACGAATTCTATCAGTTTTGGATCAACACCGACGATC
GTGACGTTATTCGTTACCTGAAGTATTTCACCTTTCTGAGCAAAGAGGAAATCGAAG
CGCTGGAGCAGGAACTGCGTGAGGCGCCGGAAAAGCGTGCGGCGCAAAAAGCGCT
GGCGGAGGAAGTGACCAAACTGGTTCACGGTGAGGAAGCGCTGCGTCAGGCGATCC
GTATTAGCGAAGCGCTGTTTAGCGGTGATATCGCGAACCTGACCGCGGCGGAGATT
GAACAAGGCTTCAAGGACGTGCCGAGCTTTGTTCACGAAGGTGGCGATGTGCCGCT
GGTTGAGCTGCTGGTTAGCGCGGGTATCAGCCCGAGCAAACGTCAGGCGCGTGAAG
ACATCCAAAACGGTGCGATTTACGTGAACGGCGAGCGTCTGCAAGATGTTGGCGCG

ATTCTGACCGCGGAACACCGTCTGGAAGGTCGTTTTACCGTTATCCGTCGTGGCAAG
AAGAAATACTATCTGATTCGTTATGCGTAA

pBK MCS Chimera H2-BPA-RS
Same as Chimera H2-YRS but with the following active site mutations:
Y37G, D179G, L183A

pBK MCS Chimera H6-YRS
Same as above but with the following aaRS sequence:
atgGCAAGCAGTAACTTGATTAAACAATTGCAAGAGCGGGGGCTGGTAGCCCAGGTG
ACGGACGAGGAAGCGTTAGCAGAGCGACTGGCGCAAGGCCCGATCGCGCTCTATTG
CGGCTTCGATCCTACCGCTGACAGCTTGCATTTGGGGCATCTTGTTCCATTGTTATGC
CTGAAACGCTTCCAGCAGGCGGGCCACAAGCCGGTTGCGCTGGTAGGCGGCGCGAC
GGGTCTGATTGGCGACCCGAGTTTCAAAGCTGCCGAGCGTAAGCTGAACACCGAAG
AAACTGTTCAGGAGTGGGTGGACAAAATCCGTAAGCAGGTTGCCCCGTTCCTCGATT
TCGACTGTGGAGAAAACTCTGCTATCGCGGCGAACAACTATGACTGGTTCGGCAATA
TGAATGTGCTGACCTTCCTGCGCGATATTGGCAAACACTTCTCCGTTAACCAGATGA
TCAACAAAGAAGCGGTTAAGCAGCGTCTCAACCGTGAAGATCAGGGGATTTCGTTC
ACTGAGTTTTCCTACAGCCTGTTGCAGGGTTATGACTTCGCCTGTCTGAACAAACAG
TACGGTGTGGTGCTGCAAATTGGTGGTTCTGACCAGTGGGGTAACATCACTTCTGGT
ATCGACCTGACCCGTCGTCTGCATCAGAATCAGGTGTTTGGCCTGACCGTTCCGCTG
ATCACTAAAGCAGATGGCACCAAATTTGGCAAAACCGAAAGCGGTACCATTTGGCT
GGATAAGGAGAAAACCAGCCCGTACGAATTCTATCAGTTTTGGATCAACACCGACG
ATCGTGACGTTATTCGTTACCTGAAGTATTTCACCTTTCTGAGCAAAGAGGAAATCG
AAGCGCTGGAGCAGGAACTGCGTGAGGCGCCGGAAAAGCGTGCGGCGCAAAAAGC
GCTGGCGGAGGAAGTGACCAAACTGGTTCACGGTGAGGAAGCGCTGCGTCAGGCGA
TCCGTATTAGCGAAGCGCTGTTTAGCGGTGATATCGCGAACCTGACCGCGGCGGAG
ATTGAACAAGGCTTCAAGGACGTGCCGAGCTTTGTTCACGAAGGTGGCGATGTGCC
GCTGGTTGAGCTGCTGGTTAGCGCGGGTATCAGCCCGAGCAAACGTCAGGCGCGTG
AAGACATCCAAAACGGTGCGATTTACGTGAACGGCGAGCGTCTGCAAGATGTTGGC
GCGATTCTGACCGCGGAACACCGTCTGGAAGGTCGTTTTACCGTTATCCGTCGTGGC
AAGAAGAAATACTATCTGATTCGTTATGCGTAA

pBK MCS Chimera H6-BPA-RS
Same as Chimera H6-YRS but with the following active site mutations:
Y37G, D182G, L186A

pET22b 10X N-term His GsYRS

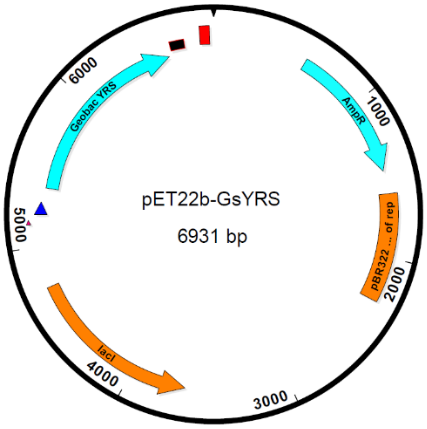


tggcgaatgggacgcgccctgtagcggcgcattaagcgcggcgggtgtggtggttacgcgcagcgtgaccgctacacttgccagcgccc
tagcgcccgctcctttcgctttcttcccttcctttctcgccacgttcgccggctttccccgtcaagctctaaatcgggggctccctttagggttcc
gatttagtgctttacggcacctcgaccccaaaaaacttgattagggtgatggttcacgtagtgggccatcgccctgatagacggtttttcgccc
tttgacgttggagtccacgttctttaatagtggactcttgttccaaactggaacaacactcaaccctatctcggtctattcttttgatttataagggat
tttgccgatttcggcctattggttaaaaaatgagctgatttaacaaaaatttaacgcgaattttaacaaaatattaacgtttacaatttcaggtggca
cttttcggggaaatgtgcgcggaacccctatttgtttatttttctaaatacattcaaatatgtatccgctcatgagacaataaccctgataaatgctt
caataatattgaaaaaggaagagtatgagtattcaacatttccgtgtcgcccttattcccttttttgcggcattttgccttcctgtttttgctcaccca
gaaacgctggtgaaagtaaaagatgctgaagatcagttgggtgcacgagtgggttacatcgaactggatctcaacagcggtaagatccttg
agagttttcgccccgaagaacgttttccaatgatgagcacttttaaagttctgctatgtggcgcggtattatcccgtattgacgccgggcaaga
gcaactcggtcgccgcatacactattctcagaatgacttggttgagtactcaccagtcacagaaaagcatcttacggatggcatgacagtaag
agaattatgcagtgctgccataaccatgagtgataacactgcggccaacttacttctgacaacgatcggaggaccgaaggagctaaccgctt
ttttgcacaacatgggggatcatgtaactcgccttgatcgttgggaaccggagctgaatgaagccataccaaacgacgagcgtgacaccac
gatgcctgcagcaatggcaacaacgttgcgcaaactattaactggcgaactacttactctagcttcccggcaacaattaatagactggatgga
ggcggataaagttgcaggaccacttctgcgctcggcccttccggctggctggtttattgctgataaatctggagccggtgagcgtgggtctc
gcggtatcattgcagcactggggccagatggtaagccctcccgtatcgtagttatctacacgacggggagtcaggcaactatggatgaacg
aaatagacagatcgctgagataggtgcctcactgattaagcattggtaactgtcagaccaagtttactcatatatactttagattgatttaaaactt
catttttaatttaaaaggatctaggtgaagatcctttttgataatctcatgaccaaaatcccttaacgtgagttttcgttccactgagcgtcagaccc
cgtagaaaagatcaaaggatcttcttgagatcctttttttctgcgcgtaatctgctgcttgcaaacaaaaaaaccaccgctaccagcggtggttt
gtttgccggatcaagagctaccaactctttttccgaaggtaactggcttcagcagagcgcagataccaaatactgtccttctagtgtagccgta
gttaggccaccacttcaagaactctgtagcaccgcctacatacctcgctctgctaatcctgttaccagtggctgctgccagtggcgataagtc
gtgtcttaccgggttggactcaagacgatagttaccggataaggcgcagcggtcgggctgaacggggggttcgtgcacacagcccagctt
ggagcgaacgacctacaccgaactgagatacctacagcgtgagctatgagaaagcgccacgcttcccgaagggagaaaggcggacag
gtatccggtaagcggcagggtcggaacaggagagcgcacgagggagcttccagggggaaacgcctggtatctttatagtcctgtcgggtt
tcgccacctctgacttgagcgtcgatttttgtgatgctcgtcaggggggcggagcctatggaaaaacgccagcaacgcggcctttttacggtt
cctggccttttgctggccttttgctcacatgttctttcctgcgttatcccctgattctgtggataaccgtattaccgcctttgagtgagctgataccg
ctcgccgcagccgaacgaccgagcgcagcgagtcagtgagcgaggaagcggaagagcgcctgatgcggtattttctccttacgcatctgt
gcggtatttcacaccgcatatatggtgcactctcagtacaatctgctctgatgccgcatagttaagccagtatacactccgctatcgctacgtga
ctgggtcatggctgcgccccgacacccgccaacacccgctgacgcgccctgacgggcttgtctgctcccggcatccgcttacagacaagc
tgtgaccgtctccgggagctgcatgtgtcagaggttttcaccgtcatcaccgaaacgcgcgaggcagctgcggtaaagctcatcagcgtgg

tcgtgaagcgattcacagatgtctgcctgttcatccgcgtccagctcgttgagtttctccagaagcgttaatgtctggcttctgataaagcgggc
catgttaagggcggtttttttcctgtttggtcactgatgcctccgtgtaagggggatttctgttcatgggggtaatgataccgatgaaacgagaga
ggatgctcacgatacgggttactgatgatgaacatgcccggttactggaacgttgtgagggtaaacaactggcggtatggatgcggcggga
ccagagaaaaatcactcagggtcaatgccagcgcttcgttaatacagatgtaggtgttccacagggtagccagcagcatcctgcgatgcag
atccggaacataatggtgcagggcgctgacttccgcgtttccagactttacgaaacacggaaaccgaagaccattcatgttgttgctcaggtc
gcagacgttttgcagcagcagtcgcttcacgttcgctcgcgtatcggtgattcattctgctaaccagtaaggcaaccccgccagcctagccg
ggtcctcaacgacaggagcacgatcatgcgcacccgtggggccgccatgccggcgataatggcctgcttctcgccgaaacgtttggtggc
gggaccagtgacgaaggcttgagcgagggcgtgcaagattccgaataccgcaagcgacaggccgatcatcgtcgcgctccagcgaaag
cggtcctcgccgaaaatgacccagagcgctgccggcacctgtcctacgagttgcatgataaagaagacagtcataagtgcggcgacgata
gtcatgccccgcgcccaccggaaggagctgactgggttgaaggctctcaagggcatcggtcgagatcccggtgcctaatgagtgagctaa
cttacattaattgcgttgcgcgtggtgaatgtgaaaccagtaacgttatacgatgtcgcagagtatgccggtgtctcttatcagaccgtttcccg
cgtggtgaaccaggccagccacgtttctgcgaaaacgcgggaaaaagtggaagcggcgatggcggagctgaattacattcccaaccgcg
tggcacaacaactggcgggcaaacagtcgttgctgattggcgttgccacctccagtctggccctgcacgcgccgtcgcaaattgtcgcggc
gattaaatctcgcgccgatcaactgggtgccagcgtggtggtgtcgatggtagaacgaagcggcgtcgaagcctgtaaagcggcggtgca
caatcttctcgcgcaacgcgtcagtgggctgatcattaactatccgctggatgaccaggatgccattgctgtggaagctgcctgcactaatgtt
ccggcgttatttcttgatgtctctgaccagacacccatcaacagtattattttctcccatgaagacggtacgcgactgggcgtggagcatctggt
cgcattgggtcaccagcaaatcgcgctgttagcgggcccattaagttctgtctcggcgcgtctgcgtctggctggctggcataaatatctcac
tcgcaatcaaattcagccgatagcggaacgggaaggcgactggagtgccatgtccggttttcaacaaaccatgcaaatgctgaatgaggg
catcgttcccactgcgatgctggttgccaacgatcagatggcgctgggcgcaatgcgcgccattaccgagtccgggctgcgcgttggtgcg
gatatctcggtagtgggatacgacgataccgaagacagctcatgttatatcccgccgttaaccaccatcaaacaggattttcgcctgctgggg
caaaccagcgtggaccgcttgctgcaactctctcagggccaggcggtgaagggcaatcagctgttgcccgtctcactggtgaaaagaaaa
accaccctggcgcccaatacgcaaaccgcctctccccgcgcgttggccgattcattaatgcagctggcacgacaggtttcccgactggaaa
gcgggcagtgacctgaattgactctcttccgggcgctatcatgccataccgcgaaaggttttgcgccattcgatggtgtccgggatctcgac
gctctcccttatgcgactcctgcattaggaagcagcccagtagtaggttgaggccgttgagcaccgccgccgcaaggaatggtgcatgcaa
ggagatggcgcccaacagtcccccggccacggggcctgccaccatacccacgccgaaacaagcgctcatgagcccgaagtggcgagc
ccgatcttccccatcggtgatgtcggcgatataggcgccagcaaccgcacctgtggcgccggtgatgccggccacgatgcgtccggcgta
gaggatcgagatctcgatcccgcgaaattaatacgactcactataggggaattgtgagcggataacaattcccctctagagtttgacagcatt
atcatcgatctcgagaaatcataaaaaatttatttgctttgtgagcggataacaattataatagattcaattgtgagcggataacaatttcacacag
aattcattaaagaggagaaattacatatgcatcatcaccatcaccatcatcatcatcacgcaagcagtaacttgattaaacaattgcaagagG
ACCTGCTGGCGGAACTGCAATGGCGTGGCCTGGTTAATCAGACCACCGACGAAGAT
GGCCTGCGTAAACTGCTGAACGAGGAACGTGTGACCCTGTATTGCGGTTTCGACCCG
ACCGCGGATAGCCTGCACATCGGCAACCTGGCGGCGATTCTGACCCTGCGTCGTTTT
CAGCAAGCGGGTCACCGTCCGATCGCGCTGGTTGGTGGTGCGACCGGTCTGATTGGC
GACCCGAGCGGCAAGAAAAGCGAGCGTACCCTGAACGCGAAGGAAACCGTTGAAG
CGTGGAGCGCGCGTATCAAAGAACAGCTGGGTCGTTTCCTGGACTTTGAGGCGGAT
GGCAACCCGGCGAAGATTAAAAACAACTATGACTGGATCGGTCCGCTGGATGTGAT
TACCTTCCTGCGTGATGTGGGCAAGCACTTTAGCGTTAACTACATGATGGCGAAAGA
GAGCGTTCAGAGCCGTATCGAAACCGGTATTAGCTTCACCGAGTTTAGCTACATGAT
GCTGCAAGCGTATGACTTCCTGCGTCTGTACGAAACCGAAGGCTGCCGTCTGCAGAT
CGGTGGCAGCGATCAATGGGGTAACATCACCGCGGGCCTGGAACTGATTCGTAAGA
CCAAAGGTGAAGCGCGTGCGTTTGGCCTGACCATCCCGCTGGTGACCAAGGCGGAC
GGTACCAAGTTTGGCAAAACCGAAAGCGGTACCATTTGGCTGGATAAGGAGAAAAC
CAGCCCGTACGAATTCTATCAGTTTTGGATCAACACCGACGATCGTGACGTTATTCG

TTACCTGAAGTATTTCACCTTTCTGAGCAAAGAGGAAATCGAAGCGCTGGAGCAGG
AACTGCGTGAGGCGCCGGAAAAGCGTGCGGCGCAAAAAGCGCTGGCGGAGGAAGT
GACCAAACTGGTTCACGGTGAGGAAGCGCTGCGTCAGGCGATCCGTATTAGCGAAG
CGCTGTTTAGCGGTGATATCGCGAACCTGACCGCGGCGGAGATTGAACAAGGCTTC
AAGGACGTGCCGAGCTTTGTTCACGAAGGTGGCGATGTGCCGCTGGTTGAGCTGCTG
GTTAGCGCGGGTATCAGCCCGAGCAAACGTCAGGCGCGTGAAGACATCCAAAACGG
TGCGATTTACGTGAACGGCGAGCGTCTGCAAGATGTTGGCGCGATTCTGACCGCGGA
ACACCGTCTGGAAGGTCGTTTTACCGTTATCCGTCGTGGCAAGAAGAAATACTATCT
GATTCGTTATGCGTAAaagcttaattagctgagcttggactcctgttgatagatccagtaatgacctcagaactccatctggattt
gttcagaacgctcggttgccgccgggcgtttttttattggtgagaatccaagctagcttggcggcggccgcactcgagcaccaccaccacca
ccactgagatccggctgctaacaaagcccgaaaggaagctgagttggctgctgccaccgctgagcaataactagcataaccccttggggc
ctctaaacgggtcttgaggggttttttgctgaaaggaggaactatatccggat

pET22b 10X N-term His Gs-BPA-RS
Same as GsYRS wt but with the following active site mutations:
Y34G, D176G, L180A

pET22b 10X N-term His MjYRS wt
Same as above but with the following aaRS sequence:
gacgaatttgaaatgataaagagaaacacatctgaaattatcagcgaggaagagttaagagaggttttaaaaaaagatgaaaaatctgctnn
nataggttttgaaccaagtggtaaaatacatttagggcattatctccaaataaaaaagatgattgatttacaaaatgctggatttgatataattata
nnnttggctgatttannngcctatttaaaccagaaaggagagttggatgagattagaaaaataggagattataacaaaaaagtttttgaagca
atggggttaaaggcaaaatatgtttatggaagtgaannnnnncttgataaggattatacactgaatgtctatagattggctttaaaaactacctt
aaaaagagcaagaaggagtatggaacttatagcaagagaggatgaaaatccaaaggttgctgaagttatctatccaataatgnnngttaatn
nnnnncattatnnnggcgttgatgttgcagttggagggatggagcagagaaaaatacacatgttagcaagggagcttttaccaaaaaaggt
tgtttgtattcacaaccctgtcttaacgggtttggatggagaaggaaagatgagttcttcaaaagggaattttatagctgttgatgactctccaga
agagattagggctaagataaagaaagcatactgcccagctggagttgttgaaggaaatccaataatggagatagctaaatacttccttgaata
tcctttaaccataaaaaggccagaaaaatttggtggagatttgacagttaatagctatgaggagttagagagtttatttaaaaataaggaattgc
atccaatggatttaaaaaatgctgtagctgaagaacttataaagattttagagccaattagaaagagattataa

pET22b 10X N-term His Mj-CNF-RS
Same as MjYRS wt but with the following active site mutations:

pET22b 10X N-term His EcYRS wt
Same as above but with the following aaRS sequence:
GCAAGCAGTAACTTGATTAAACAATTGCAAGAGCGGGGGCTGGTAGCCCAGGTGAC
GGACGAGGAAGCGTTAGCAGAGCGACTGGCGCAAGGCCCGATCGCGCTCTATTGCG
GCTTCGATCCTACCGCTGACAGCTTGCATTTGGGGCATCTTGTTCCATTGTTATGCCT
GAAACGCTTCCAGCAGGCGGGCCACAAGCCGGTTGCGCTGGTAGGCGGCGCGACGG
GTCTGATTGGCGACCCGAGCTTCAAAGCTGCCGAGCGTAAGCTGAACACCGAAGAA
ACTGTTCAGGAGTGGGTGGACAAAATCCGTAAGCAGGTTGCCCCGTTCCTCGATTTC
GACTGTGGAGAAAACTCTGCTATCGCGGCGAACAACTATGACTGGTTCGGCAATAT
GAATGTGCTGACCTTCCTGCGCGATATTGGCAAACACTTCTCCGTTAACCAGATGAT
CAACAAAGAAGCGGTTAAGCAGCGTCTCAACCGTGAAGATCAGGGGATTTCGTTCA

CTGAGTTTTCCTACAACCTGTTGCAGGGTTATGACTTCGCaTGTCTGAACAAACAGTA
CGGTGTGGTGCTGCAAATTGGTGGTTCTGACCAGTGGGGTAACATCACTTCTGGTAT
CGACCTGACCCGTCGTCTGCATCAGAATCAGGTGTTTGGCCTGACCGTTCCGCTGAT
CACTAAAGCAGATGGCACCAAATTTGGTAAAACTGAAGGCGGCGCAGTCTGGTTaG
ATCCGAAGAAAACCAGCCCGTACAAATTCTACCAGTTCTGGATCAACACTGCGGAC
GCCGACGTTTACCGCTTCCTGAAGTTCTTCACCTTTATGAGCATTGAAGAGATCAAC
GCCCTGGAAGAAGAAGATAAAAACAGCGGTAAAGCACCGCGCGCCCAGTATGTACT
GGCGGAGCAGGTGACTCGTCTGGTTCACGGTGAAGAAGGTTTACAGGCGGCAAAAC
GTATTACCGAATGCCTGTTCAGCGGTTCTTTGAGTGCGCTGAGTGAAGCGGACTTCG
AACAGCTGGCGCAGGACGGCGTACCGATGGTTGAGATGGAAAAGGGCGCAGACCTG
ATGCAGGCACTGGTCGATTCTGAACTGCAACCTTCCCGTGGTCAGGCACGTAAAACT
ATCGCCTCCAATGCCATCACCATTAACGGTGAAAAACAGTCCGATCCTGAATACTTC
TTTAAAGAAGAAGATCGTCTGTTTGGTCGTTTTACCTTACTGCGTCGCGGTAAAAAG
AATTACTGTCTGATTTGCTGGAAAtaa

pET22b 10X N-term His Ec-BPA-RS
Same as EcYRS wt but with the following active site mutations:
Y37G, D182G, L186A

pET22b 10X N-term His Ec-OMeYRS
Same as EcYRS wt but with the following active site mutations:
Y37V, D182S, F183M, L186A

pET22b 10X N-term His Chimera H2-YRS
Same as above but with the following aaRS sequence:
GCAAGCAGTAACTTGATTAAACAATTGCAAGAGCGGGGGCTGGTAGCCCAGGTGAC
GGACGAGGAAGCGTTAGCAGAGCGACTGGCGCAAGGCCCGATCGCGCTCTATTGCG
GCTTCGATCCTACCGCTGACAGCTTGCATTTGGGGCATCTTGTTCCATTGTTATGCCT
GAAACGCTTCCAGCAGGCGGGTCACCGTCCGATCGCGCTGGTTGGTGGTGCGACCG
GTCTGATTGGCGACCCGAGCGGCAAGAAAAGCGAGCGTACCCTGAACGCGAAGGA
AACCGTTGAAGCGTGGAGCGCGCGTATCAAAGAACAGCTGGGTCGTTTCCTGGACTT
TGAGGCGGATGGCAACCCGGCGAAGATTAAAAACAACTATGACTGGATCGGTCCGC
TGGATGTGATTACCTTCCTGCGTGATGTGGGCAAGCACTTTAGCGTTAACTACATGA
TGGCGAAAGAGAGCGTTCAGAGCCGTATCGAAACCGGTATTAGCTTCACCGAGTTT
AGCTACATGATGCTGCAAGCGTATGATTTCCTGCGTCTGTACGAAACCGAAGGCTGC
CGTCTGCAGATCGGTGGCAGCGATCAATGGGGTAACATCACCGCGGGCCTGGAACT
GATTCGTAAGACCAAAGGTGAAGCGCGTGCGTTTGGCCTGACCATCCCGCTGGTGAC
CAAGGCGGACGGTACCAAGTTTGGCAAAACCGAAAGCGGTACCATTTGGCTGGATA
AGGAGAAAACCAGCCCGTACGAATTCTATCAGTTTTGGATCAACACCGACGATCGT
GACGTTATTCGTTACCTGAAGTATTTCACCTTTCTGAGCAAAGAGGAAATCGAAGCG
CTGGAGCAGGAACTGCGTGAGGCGCCGGAAAAGCGTGCGGCGCAAAAAGCGCTGG
CGGAGGAAGTGACCAAACTGGTTCACGGTGAGGAAGCGCTGCGTCAGGCGATCCGT
ATTAGCGAAGCGCTGTTTAGCGGTGATATCGCGAACCTGACCGCGGCGGAGATTGA
ACAAGGCTTCAAGGACGTGCCGAGCTTTGTTCACGAAGGTGGCGATGTGCCGCTGGT

TGAGCTGCTGGTTAGCGCGGGTATCAGCCCGAGCAAACGTCAGGCGCGTGAAGACA
TCCAAAACGGTGCGATTTACGTGAACGGCGAGCGTCTGCAAGATGTTGGCGCGATTC
TGACCGCGGAACACCGTCTGGAAGGTCGTTTTACCGTTATCCGTCGTGGCAAGAAGA
AATACTATCTGATTCGTTATGCGTAA

pET 22b 10X N-term His Chimera H2-BPA-RS
Same as H2-YRS but with the following active site mutations:
Y37G, D179G, L183A

pET22b 10X N-term His Chimera H6-YRS
Same as above but with the following aaRS sequence:
GCAAGCAGTAACTTGATTAAACAATTGCAAGAGCGGGGGCTGGTAGCCCAGGTGAC
GGACGAGGAAGCGTTAGCAGAGCGACTGGCGCAAGGCCCGATCGCGCTCTATTGCG
GCTTCGATCCTACCGCTGACAGCTTGCATTTGGGGCATCTTGTTCCATTGTTATGCCT
GAAACGCTTCCAGCAGGCGGGCCACAAGCCGGTTGCGCTGGTAGGCGGCGCGACGG
GTCTGATTGGCGACCCGAGTTTCAAAGCTGCCGAGCGTAAGCTGAACACCGAAGAA
ACTGTTCAGGAGTGGGTGGACAAAATCCGTAAGCAGGTTGCCCCGTTCCTCGATTTC
GACTGTGGAGAAAACTCTGCTATCGCGGCGAACAACTATGACTGGTTCGGCAATAT
GAATGTGCTGACCTTCCTGCGCGATATTGGCAAACACTTCTCCGTTAACCAGATGAT
CAACAAAGAAGCGGTTAAGCAGCGTCTCAACCGTGAAGATCAGGGGATTTCGTTCA
CTGAGTTTTCCTACAGCCTGTTGCAGGGTTATGACTTCGCCTGTCTGAACAAACAGT
ACGGTGTGGTGCTGCAAATTGGTGGTTCTGACCAGTGGGGTAACATCACTTCTGGTA
TCGACCTGACCCGTCGTCTGCATCAGAATCAGGTGTTTGGCCTGACCGTTCCGCTGA
TCACTAAAGCAGATGGCACCAAATTTGGCAAAACCGAAAGCGGTACCATTTGGCTG
GATAAGGAGAAAACCAGCCCGTACGAATTCTATCAGTTTTGGATCAACACCGACGA
TCGTGACGTTATTCGTTACCTGAAGTATTTCACCTTTCTGAGCAAAGAGGAAATCGA
AGCGCTGGAGCAGGAACTGCGTGAGGCGCCGGAAAAGCGTGCGGCGCAAAAAGCG
CTGGCGGAGGAAGTGACCAAACTGGTTCACGGTGAGGAAGCGCTGCGTCAGGCGAT
CCGTATTAGCGAAGCGCTGTTTAGCGGTGATATCGCGAACCTGACCGCGGCGGAGA
TTGAACAAGGCTTCAAGGACGTGCCGAGCTTTGTTCACGAAGGTGGCGATGTGCCGC
TGGTTGAGCTGCTGGTTAGCGCGGGTATCAGCCCGAGCAAACGTCAGGCGCGTGAA
GACATCCAAAACGGTGCGATTTACGTGAACGGCGAGCGTCTGCAAGATGTTGGCGC
GATTCTGACCGCGGAACACCGTCTGGAAGGTCGTTTTACCGTTATCCGTCGTGGCAA
GAAGAAATACTATCTGATTCGTTATGCGTAA

pET22b 10X N-term His Chimera H6-BPA-RS
Same as H6-YRS but with the following active site mutations:
Y37G, D182G, L186A

pB1U Gs-BPA-RS

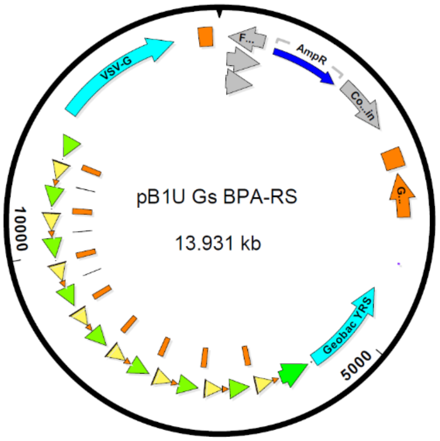


ttctctgtcacagaatgaaaatttttctgtcatctcttcgttattaatgtttgtaattgactgaatatcaacgcttatttgcagcctgaatggcgaatgg
gacgcgccctgtagcggcgcattaagcgcggcgggtgtggtggttacgcgcagcgtgaccgctacacttgccagcgccctagcgcccgc
tcctttcgctttcttcccttcctttctcgccacgttcgccggctttccccgtcaagctctaaatcgggggctccctttagggttccgatttagtgcttt
acggcacctcgaccccaaaaaacttgattagggtgatggttcacgtagtgggccatcgccctgatagacggtttttcgccctttgacgttgga
gtccacgttctttaatagtggactcttgttccaaactggaacaacactcaaccctatctcggtctattcttttgatttataagggattttgccgatttc
ggcctattggttaaaaaatgagctgatttaacaaaaatttaacgcgaattttaacaaaatattaacgtttacaatttcaggtggcacttttcgggga
aatgtgcgcggaacccctatttgtttatttttctaaatacattcaaatatgtatccgctcatgagacaataaccctgataaatgcttcaataatattg
aaaaaggaagagtatgagtattcaacatttccgtgtcgcccttattcccttttttgcggcattttgccttcctgtttttgctcacccagaaacgctgg
tgaaagtaaaagatgctgaagatcagttgggtgcacgagtgggttacatcgaactggatctcaacagcggtaagatccttgagagttttcgcc
ccgaagaacgttttccaatgatgagcacttttaaagttctgctatgtggcgcggtattatcccgtattgacgccgggcaagagcaactcggtcg
ccgcatacactattctcagaatgacttggttgagtactcaccagtcacagaaaagcatcttacggatggcatgacagtaagagaattatgcag
tgctgccataaccatgagtgataacactgcggccaacttacttctgacaacgatcggaggaccgaaggagctaaccgcttttttgcacaacat
gggggatcatgtaactcgccttgatcgttgggaaccggagctgaatgaagccataccaaacgacgagcgtgacaccacgatgcctgtagc
aatggcaacaacgttgcgcaaactattaactggcgaactacttactctagcttcccggcaacaattaatagactggatggaggcggataaagt
tgcaggaccacttctgcgctcggcccttccggctggctggtttattgctgataaatctggagccggtgagcgtgggtctcgcggtatcattgc
agcactggggccagatggtaagccctcccgtatcgtagttatctacacgacggggagtcaggcaactatggatgaacgaaatagacagat
cgctgagataggtgcctcactgattaagcattggtaactgtcagaccaagtttactcatatatactttagattgatttaaaacttcatttttaatttaa
aaggatctaggtgaagatcctttttgataatctcatgaccaaaatcccttaacgtgagttttcgttccactgagcgtcagaccccgtagaaaaga
tcaaaggatcttcttgagatcctttttttctgcgcgtaatctgctgcttgcaaacaaaaaaaccaccgctaccagcggtggtttgtttgccggatc
aagagctaccaactctttttccgaaggtaactggcttcagcagagcgcagataccaaatactgtccttctagtgtagccgtagttaggccacc
acttcaagaactctgtagcaccgcctacatacctcgctctgctaatcctgttaccagtggctgctgccagtggcgataagtcgtgtcttaccgg
gttggactcaagacgatagttaccggataaggcgcagcggtcgggctgaacggggggttcgtgcacacagcccagcttggagcgaacg
acctacaccgaactgagatacctacagcgtgagcattgagaaagcgccacgcttcccgaagggagaaaggcggacaggtatccggtaag
cggcagggtcggaacaggagagcgcacgagggagcttccagggggaaacgcctggtatctttatagtcctgtcgggtttcgccacctctg
acttgagcgtcgatttttgtgatgctcgtcaggggggcggagcctatggaaaaacgccagcaacgcggcctttttacggttcctggccttttg
ctggccttttgctcacatgttctttcctgcgttatcccctgattctgtggataaccgtattaccgcctttgagtgagctgataccgctcgccgcagc
cgaacgaccgagcgcagcgagtcagtgagcgaggaagcggaagagcgcctgatgcggtattttctccttacgcatctgtgcggtatttcac
accgcagaccagccgcgtaacctggcaaaatcggttacggttgagtaataaatggatgccctgcgtaagcgggtgtgggcggacaataaa
gtcttaaactgaacaaaatagatctaaactatgacaataaagtcttaaactagacagaatagttgtaaactgaaatcagtccagttatgctgtga
aaaagcatactggacttttgttatggctaaagcaaactcttcattttctgaagtgcaaattgcccgtcgtattaaagaggggcgtggccaaggg
catggtaaagactatattcgcggcgttgtgacaatttaccgaacaactccgcggccgggaagccgatctcggcttgaacgaattgatgttac

gcagcagcaacgatgttacgcagcagggcagtcgccctaaaacaaagttaggtggctcaagtatgggcatcattcgcacatgtaggctcg
gccctgaccaagtcaaatccatgcgggctgctcttgatcttttcggtcgtgagttcggagacgtagccacctactcccaacatcagccggact
ccgattacctcgggaacttgctccgtagtaagacattcatcgcgcttgctgccttcgaccaagaagcggttgttggcgctctcgcggcttacgt
tctgcccaggtttgagcagccgcgtagtgagatctatatctatgatctcgcagtctccggcgagcaccggaggcagggcattgccaccgcg
ctcatcaatctcctcaagcatgaggccaacgcgcttggtgcttatgtgatctacgtgcaagcagattacggtgacgatcccgcagtggctctc
tatacaaagttgggcatacgggaagaagtgatgcactttgatatcgacccaagtaccgccacctaacgttgctgctccataacatcaaacatc
gacccacggcgtaacgcgcttgctgcttggatgcccgaggcatagactgtacaaaaaaacagtcataacaagccatgaaaaccgccactg
cgccgttaccaccgctgcgttcggtcaaggttctggaccagttgcgtgagcgcatacgctacttgcattacagtttacgaaccgaacaggctt
atgtcaactgggttcgtgccttcatccgtttccacggtgtgcgtcacccggcaaccttgggcagcagcgaagtcgaggcatttctgtcctggc
tggcgaacgagcgcaaggtttcggtctccacgcatcgtcaggcattggcggccttgctgttcttctacggcaaggtgctgtgcacggatctg
ccctggcttcaggagatcggtagacctcggccgtcgcggcgcttgccggtggtgctgaccccggatgaagtggttcgcatcctcggttttct
ggaaggcgagcatcgtttgttcgcccaggactctagctatagttctagtggttggctacgtacccgtagtggctatggcagggcttgcgctta
atgcgccgctacagggcgcgtggggataccccctagagccccagctggttctttccgcctcagaagccatagagcccaccgcatccccag
catgcctgctattgtcttcccaatcctcccccttgctgtcctgccccaccccaccccccagaatagaatgacacctactcagacaatgcgatgc
aatttcctcattttattaggaaaggacagtgggagtggcaccttccagggtcaaggaaggcacgggggaggggcaaacaacagatggctg
gcaactagaaggcacagtcgaggctgatcagcgggtttaaacgggccctctagactcgagttaaagtcgacttaacgcgttgaattcTTA
CGCATAACGAATCAGATAGTATTTCTTCTTGCCACGACGGATAACGGTAAAACGACC
TTCCAGACGGTGTTCCGCGGTCAGAATCGCGCCAACATCTTGCAGACGCTCGCCGTT
CACGTAAATCGCACCGTTTTGGATGTCTTCACGCGCCTGACGTTTGCTCGGGCTGAT
ACCCGCGCTAACCAGCAGCTCAACCAGCGGCACATCGCCACCTTCGTGAACAAAGC
TCGGCACGTCCTTGAAGCCTTGTTCAATCTCCGCCGCGGTCAGGTTCGCGATATCAC
CGCTAAACAGCGCTTCGCTAATACGGATCGCCTGACGCAGCGCTTCCTCACCGTGAA
CCAGTTTGGTCACTTCCTCCGCCAGCGCTTTTTGCGCCGCACGCTTTTCCGGCGCCTC
ACGCAGTTCCTGCTCCAGCGCTTCGATTTCCTCTTTGCTCAGAAAGGTGAAATACTTC
AGGTAACGAATAACGTCACGATCGTCGGTGTTGATCCAAAACTGATAGAATTCGTAC
GGGCTGGTTTTCTCCTTATCCAGCCAAATGGTACCGCTTTCGGTTTTGCCAAACTTGG
TACCGTCCGCCTTGGTCACCAGCGGGATGGTCAGGCCAAACGCACGCGCTTCACCTT
TGGTCTTACGAATCAGTTCCAGGCCCGCGGTGATGTTACCCCATTGATCGCTGCCAC
CGATCTGCAGACGGCAGCCTTCGGTTTCGTAGGCACGCAGGAAACCATACGCTTGC
AGCATCATGTAGCTAAACTCGGTGAAGCTAATACCGGTTTCGATACGGCTCTGAACG
CTCTCTTTCGCCATCATGTAGTTAACGCTAAAGTGCTTGCCCACATCACGCAGGAAG
GTAATCACATCCAGCGGACCGATCCAGTCATAGTTGTTTTTAATCTTCGCCGGGTTG
CCATCCGCCTCAAAGTCCAGGAAACGACCCAGCTGTTCTTTGATACGCGCGCTCCAC
GCTTCAACGGTTTCCTTCGCGTTCAGGGTACGCTCGCTTTTCTTGCCGCTCGGGTCGC
CAATCAGACCGGTCGCACCACCAACCAGCGCGATCGGACGGTGACCCGCTTGCTGA
AAACGACGCAGGGTCAGAATCGCCGCCAGGTTGCCGATGTGCAGGCTATCCGCGGT
CGGGTCGAAACCGCAACCCAGGGTCACACGTTCCTCGTTCAGCAGTTTACGCAGGCC
ATCTTCGTCGGTGGTCTGATTAACCAGGCCACGCCATTGCAGTTCCGCCAGCAGGTC
CATggtggcgctagccagcttgggtctccctatagtgagtcgtattaatttcgataagccagtaagcagtgggttctctagttagccagaga
gctCtagaccaagtgacgatcacagcgatccacaaacaagaaccgcgacccaaatcccggctgcgacggaactagctgtgccacaccc
ggcgcgtccttatataatcatcggcgttcaccgccccacggagatccctccgcagaatcgccgagaagggactacttttcctcgcctgttcc
gctctctggaaagaaaaccagtgccctagagtcacccaagtcccgtcctaaaatgtccttctgctgatactggggttctaaggccgagtcttat
gagcagcgggccgctgtcctgagcgtccgggcggaaggatcaggacgctcgctgcgcccttcgtctgacgtggcagcgctcgccgtga

ggagggggcgcccgcgggaggcgccaaaacccggcgcggaggccttcgaacggCCActagcaaaaaaTGGAgGGGGA
CGGATTCGAACCGCCGAACCCAAAGGGAGCGGATTtgaAGTCCGCCGCGTTTAGCCA
CTTCGCTACCCCTCCgggGATCTGTGGTCTCATACAGAACTTATAAGATTCCCAAATC
CAAAGACATTTCACGTTTATGGTGATTTCCCAGAACACATAGCGACATGCAAATATT
GCAGGGCGCCACTCCCCTGTCCCTCACAGCCATCTTCCTGCCAGGGCGCACGCGCGC
TGGGTGTTCCCGCCTAGTGACACTGGGCCCGCGATTCCTTGGAGCGGGTTGATGACG
TCAGCGTTCGAATTCCTAGCAAAAAATGGTGGGGGAAGGATTCGAACCTTCGAAGT
CTGTGACGGCAGATTTgaAGTCTGCTCCCTTTGGCCGCTCGGGAACCCCACCGGTGTT
TCGTCCTTTCCACAAGATATATAAAGCCAAGAAATCGAAATACTTTCAAGTTACGGT
AAGCATATGATAGTCCATTTTAAAACATAATTTTAAAACTGCAAACTACCCAAGAAA
TTATTACTTTCTACGTCACGTATTTTGTACTAATATCTTTGTGTTTACAGTCAAATTAA
TTCTAATTATCTCTCTAACAGCCTTGTATCGTATATGCAAATATGAAGGAATCATGG
GAAATAGGCCCTCTTCCTGCCCGACCTAGcaaaaaaTGGAgGGGGACGGATTCGAACCG
CCGAACCCAAAGGGAGCGGATTtgaAGTCCGCCGCGTTTAGCCACTTCGCTACCCCTC
CgggGATCTGTGGTCTCATACAGAACTTATAAGATTCCCAAATCCAAAGACATTTCAC
GTTTATGGTGATTTCCCAGAACACATAGCGACATGCAAATATTGCAGGGCGCCACTC
CCCTGTCCCTCACAGCCATCTTCCTGCCAGGGCGCACGCGCGCTGGGTGTTCCCGCC
TAGTGACACTGGGCCCGCGATTCCTTGGAGCGGGTTGATGACGTCAGCGTTCGAATT
CCTAGCAAAAAATGGTGGGGGAAGGATTCGAACCTTCGAAGTCTGTGACGGCAGAT
TTgaAGTCTGCTCCCTTTGGCCGCTCGGGAACCCCACCGGTGTTTCGTCCTTTCCACA
AGATATATAAAGCCAAGAAATCGAAATACTTTCAAGTTACGGTAAGCATATGATAG
TCCATTTTAAAACATAATTTTAAAACTGCAAACTACCCAAGAAATTATTACTTTCTAC
GTCACGTATTTTGTACTAATATCTTTGTGTTTACAGTCAAATTAATTCTAATTATCTCT
CTAACAGCCTTGTATCGTATATGCAAATATGAAGGAATCATGGGAAATAGGCCCTCT
TCCTGCCCGACctagcaaaaaaTGGAgGGGGACGGATTCGAACCGCCGAACCCAAAGGGA
GCGGATTtgaAGTCCGCCGCGTTTAGCCACTTCGCTACCCCTCCgggGATCTGTGGTCTC
ATACAGAACTTATAAGATTCCCAAATCCAAAGACATTTCACGTTTATGGTGATTTCC
CAGAACACATAGCGACATGCAAATATTGCAGGGCGCCACTCCCCTGTCCCTCACAG
CCATCTTCCTGCCAGGGCGCACGCGCGCTGGGTGTTCCCGCCTAGTGACACTGGGCC
CGCGATTCCTTGGAGCGGGTTGATGACGTCAGCGTTCGAATTCCTAGCAAAAAATGG
TGGGGGAAGGATTCGAACCTTCGAAGTCTGTGACGGCAGATTTgaAGTCTGCTCCCTT
TGGCCGCTCGGGAACCCCACCGGTGTTTCGTCCTTTCCACAAGATATATAAAGCCAA
GAAATCGAAATACTTTCAAGTTACGGTAAGCATATGATAGTCCATTTTAAAACATAA
TTTTAAAACTGCAAACTACCCAAGAAATTATTACTTTCTACGTCACGTATTTTGTACT
AATATCTTTGTGTTTACAGTCAAATTAATTCTAATTATCTCTCTAACAGCCTTGTATC
GTATATGCAAATATGAAGGAATCATGGGAAATAGGCCCTCTTCCTGCCCGACCTAGc
aaaaaaGGAGGGGTAGCGAAGTGGCTAAACGCGGCGGACTtcaAATCCGCTCCCTTTGGG
TTCGGCGGTTCGAATCCGTCCCCcTCCAgggGATCTGTGGTCTCATACAGAACTTATAA
GATTCCCAAATCCAAAGACATTTCACGTTTATGGTGATTTCCCAGAACACATAGCGA
CATGCAAATATTGCAGGGCGCCACTCCCCTGTCCCTCACAGCCATCTTCCTGCCAGG
GCGCACGCGCGCTGGGTGTTCCCGCCTAGTGACACTGGGCCCGCGATTCCTTGGAGC
GGGTTGATGACGTCAGCGTTCGAATTCCTAGCAAAAAATGGTGGGGGAAGGATTCG
AACCTTCGAAGTCTGTGACGGCAGATTTgaAGTCTGCTCCCTTTGGCCGCTCGGGAAC

CCCACCGGTGTTTCGTCCTTTCCACAAGATATATAAAGCCAAGAAATCGAAATACTT
TCAAGTTACGGTAAGCATATGATAGTCCATTTTAAAACATAATTTTAAAACTGCAAA
CTACCCAAGAAATTATTACTTTCTACGTCACGTATTTTGTACTAATATCTTTGTGTTT
ACAGTCAAATTAATTCTAATTATCTCTCTAACAGCCTTGTATCGTATATGCAAATATG
AAGGAATCATGGGAAATAGGCCCTCTTCCTGCCCGACctagcaaaaaaGGAGGGGTAGCG
AAGTGGCTAAACGCGGCGGACTtcaAATCCGCTCCCTTTGGGTTCGGCGGTTCGAATC
CGTCCCCcTCCAgggGATCTGTGGTCTCATACAGAACTTATAAGATTCCCAAATCCAA
AGACATTTCACGTTTATGGTGATTTCCCAGAACACATAGCGACATGCAAATATTGCA
GGGCGCCACTCCCCTGTCCCTCACAGCCATCTTCCTGCCAGGGCGCACGCGCGCTGG
GTGTTCCCGCCTAGTGACACTGGGCCCGCGATTCCTTGGAGCGGGTTGATGACGTCA
GCGTTCGAATTCCTAGCAAAAAATGGTGGGGGAAGGATTCGAACCTTCGAAGTCTG
TGACGGCAGATTTgaAGTCTGCTCCCTTTGGCCGCTCGGGAACCCCACCGGTGTTTCG
TCCTTTCCACAAGATATATAAAGCCAAGAAATCGAAATACTTTCAAGTTACGGTAAG
CATATGATAGTCCATTTTAAAACATAATTTTAAAACTGCAAACTACCCAAGAAATTA
TTACTTTCTACGTCACGTATTTTGTACTAATATCTTTGTGTTTACAGTCAAATTAATTC
TAATTATCTCTCTAACAGCCTTGTATCGTATATGCAAATATGAAGGAATCATGGGAA
ATAGGCCCTCTTCCTGCCCGACCTAGcaaaaaaTGGAgGGGGACGGATTCGAACCGCCG
AACCCAAAGGGAGCGGATTtgaAGTCCGCCGCGTTTAGCCACTTCGCTACCCCTCCggg
GATCTGTGGTCTCATACAGAACTTATAAGATTCCCAAATCCAAAGACATTTCACGTT
TATGGTGATTTCCCAGAACACATAGCGACATGCAAATATTGCAGGGCGCCACTCCCC
TGTCCCTCACAGCCATCTTCCTGCCAGGGCGCACGCGCGCTGGGTGTTCCCGCCTAG
TGACACTGGGCCCGCGATTCCTTGGAGCGGGTTGATGACGTCAGCGTTCGAATTCCT
AGCAAAAAATGGTGGGGGAAGGATTCGAACCTTCGAAGTCTGTGACGGCAGATTTga
AGTCTGCTCCCTTTGGCCGCTCGGGAACCCCACCGGTGTTTCGTCCTTTCCACAAGAT
ATATAAAGCCAAGAAATCGAAATACTTTCAAGTTACGGTAAGCATATGATAGTCCAT
TTTAAAACATAATTTTAAAACTGCAAACTACCCAAGAAATTATTACTTTCTACGTCA
CGTATTTTGTACTAATATCTTTGTGTTTACAGTCAAATTAATTCTAATTATCTCTCTAA
CAGCCTTGTATCGTATATGCAAATATGAAGGAATCATGGGAAATAGGCCCTCTTCCT
GCCCGACctagcaaaaaaGGAGGGGTAGCGAAGTGGCTAAACGCGGCGGACTtcaAATCCG
CTCCCTTTGGGTTCGGCGGTTCGAATCCGTCCCCcTCCAgggGATCTGTGGTCTCATAC
AGAACTTATAAGATTCCCAAATCCAAAGACATTTCACGTTTATGGTGATTTCCCAGA
ACACATAGCGACATGCAAATATTGCAGGGCGCCACTCCCCTGTCCCTCACAGCCATC
TTCCTGCCAGGGCGCACGCGCGCTGGGTGTTCCCGCCTAGTGACACTGGGCCCGCGA
TTCCTTGGAGCGGGTTGATGACGTCAGCGTTCGAATTCCTAGCAAAAAATGGTGGGG
GAAGGATTCGAACCTTCGAAGTCTGTGACGGCAGATTTgaAGTCTGCTCCCTTTGGCC
GCTCGGGAACCCCACCGGTGTTTCGTCCTTTCCACAAGATATATAAAGCCAAGAAAT
CGAAATACTTTCAAGTTACGGTAAGCATATGATAGTCCATTTTAAAACATAATTTTA
AAACTGCAAACTACCCAAGAAATTATTACTTTCTACGTCACGTATTTTGTACTAATAT
CTTTGTGTTTACAGTCAAATTAATTCTAATTATCTCTCTAACAGCCTTGTATCGTATA
TGCAAATATGAAGGAATCATGGGAAATAGGCCCTCTTCCTGCCCGACCTAGcaaaaaaG
GAGGGGTAGCGAAGTGGCTAAACGCGGCGGACTtcaAATCCGCTCCCTTTGGGTTCGG
CGGTTCGAATCCGTCCCCcTCCAgggGATCTGTGGTCTCATACAGAACTTATAAGATTC
CCAAATCCAAAGACATTTCACGTTTATGGTGATTTCCCAGAACACATAGCGACATGC

AAATATTGCAGGGCGCCACTCCCCTGTCCCTCACAGCCATCTTCCTGCCAGGGCGCA
CGCGCGCTGGGTGTTCCCGCCTAGTGACACTGGGCCCGCGATTCCTTGGAGCGGGTT
GATGACGTCAGCGTTCGAATTCCTAGCAAAAAATGGTGGGGGAAGGATTCGAACCT
TCGAAGTCTGTGACGGCAGATTTgaAGTCTGCTCCCTTTGGCCGCTCGGGAACCCCAC
CGGTGTTTCGTCCTTTCCACAAGATATATAAAGCCAAGAAATCGAAATACTTTCAAG
TTACGGTAAGCATATGATAGTCCATTTTAAAACATAATTTTAAAACTGCAAACTACC
CAAGAAATTATTACTTTCTACGTCACGTATTTTGTACTAATATCTTTGTGTTTACAGT
CAAATTAATTCTAATTATCTCTCTAACAGCCTTGTATCGTATATGCAAATATGAAGG
AATCATGGGAAATAGGCCCTCTTCCTGCCCGACctagtcaataatcaatgtcaacgcgtatatctggcccgta
catcgcgaagcagcgcaaaacGGATCCtgcaggtatttGCGGCCGCggtccgtatactccggaatattaatagatcatggagat
aattaaaatgataaccatctcgcaaataaataagtattttactgttttcgtaacagttttgtaataaaaaaacctataaatattccggattattcatac
cgtcccaccatcgggcgcgAACTCCTAAAAAACCGCCACCatgaagtgccttttgtacttagcctttttattcattggggtg
aattgcaagttcaccatagtttttccacacaaccaaaaaggaaactggaaaaatgttccttctaattaccattattgcccgtcaagctcagattta
aattggcataatgacttaataggcacagccttacaagtcaaaatgcccaagagtcacaaggctattcaagcagacggttggatgtgtcatgct
tccaaatgggtcactacttgtgatttccgctggtatggaccgaagtatataacacattccatccgatccttcactccatctgtagaacaatgcaa
ggaaagcattgaacaaacgaaacaaggaacttggctgaatccaggcttccctcctcaaagttgtggatatgcaactgtgacggatgccgaa
gcagtgattgtccaggtgactcctcaccatgtgctggttgatgaatacacaggagaatgggttgattcacagttcatcaacggaaaatgcagc
aattacatatgccccactgtccataactctacaacctggcattctgactataaggtcaaagggctatgtgattctaacctcatttccatggacatc
accttcttctcagaggacggagagctatcatccctgggaaaggagggcacagggttcagaagtaactactttgcttatgaaactggaggcaa
ggcctgcaaaatgcaatactgcaagcattggggagtcagactcccatcaggtgtctggttcgagatggctgataaggatctctttgctgcagc
cagattccctgaatgcccagaagggtcaagtatctctgctccatctcagacctcagtggatgtaagtctaattcaggacgttgagaggatcttg
gattattccctctgccaagaaacctggagcaaaatcagagcgggtcttccaatctctccagtggatctcagctatcttgctcctaaaaacccag
gaaccggtcctgctttcaccataatcaatggtaccctaaaatactttgagaccagatacatcagagtcgatattgctgctccaatcctctcaaga
atggtcggaatgatcagtggaactaccacagaaagggaactgtgggatgactgggcaccatatgaagacgtggaaattggacccaatgga
gttctgaggaccagttcaggatataagtttcctttatacatgattggacatggtatgttggactccgatcttcatcttagctcaaaggctcaggtgt
tcgaacatcctcacattcaagacgctgcttcgcaacttcctgatgatgagagtttattttttggtgatactgggctatccaaaaatccaatcgagc
ttgtagaaggttggttcagtagttggaaaagctctattgcctctttttctttatcatagggttaatcattggactattcttggttctccgagttggtatc
catctttgcattaaattaaagcacaccaagaaaagacagatttatacagacatagagatgaaccgacttggaaagtgataaggccaggccgg
ccaagcttgtcgagaagtactagaggatcataatcagccataccacatttgtagaggttttacttgctttaaaaaacctcccacacctccccctg
aacctgaaacataaaatgaatgcaattgttgttgttaacttgtttattgcagcttataatggttacaaataaagcaatagcatcacaaatttcacaa
ataaagcatttttttcactgcattctagttgtggtttgtccaaactcatcaatgtatcttatcatgtctggatctgatcactgcttgagcctaggagat
ccgaaccagataagtgaaatctagttccaaactattttgtcattttaattttcgtattagcttacgacgctacacccagttcccatctattttgtcact
cttccctaaataatccttaaaaactccatttccacccctcccagttcccaactattttgtccgcccacagcggggcatttttcttcctgttatgtttttta
atcaaacatcctgccaactccatgtgacaaaccgtcatcttcggctacttt

pB1U Ec-BPA-RS
Same as above but with the following aaRS sequence:
atgGCAAGCAGTAACTTGATTAAACAATTGCAAGAGCGGGGGCTGGTAGCCCAGGTG
ACGGACGAGGAAGCGTTAGCAGAGCGACTGGCGCAAGGCCCGATCGCGCTCGGTTG
CGGCTTCGATCCTACCGCTGACAGCTTGCATTTGGGGCATCTTGTTCCATTGTTATGC
CTGAAACGCTTCCAGCAGGCGGGCCACAAGCCGGTTGCGCTGGTAGGCGGCGCGAC
GGGTCTGATTGGCGACCCGAGCTTCAAAGCTGCCGAGCGTAAGCTGAACACCGAAG
AAACTGTTCAGGAGTGGGTGGACAAAATCCGTAAGCAGGTTGCCCCGTTCCTCGATT
TCGACTGTGGAGAAAACTCTGCTATCGCGGCGAACAACTATGACTGGTTCGGCAATA

TGAATGTGCTGACCTTCCTGCGCGATATTGGCAAACACTTCTCCGTTAACCAGATGA
TCAACAAAGAAGCGGTTAAGCAGCGTCTCAACCGTGAAGATCAGGGGATTTCGTTC
ACTGAGTTTTCCTACAACCTGTTGCAGGGTTATGGTTTCGCaTGTGCTAACAAACAGT
ACGGTGTGGTGCTGCAAATTGGTGGTTCTGACCAGTGGGGTAACATCACTTCTGGTA
TCGACCTGACCCGTCGTCTGCATCAGAATCAGGTGTTTGGCCTGACCGTTCCGCTGA
TCACTAAAGCAGATGGCACCAAATTTGGTAAAACTGAAGGCGGCGCAGTCTGGTTaG
ATCCGAAGAAAACCAGCCCGTACAAATTCTACCAGTTCTGGATCAACACTGCGCGT
GCCGACGTTTACCGCTTCCTGAAGTTCTTCACCTTTATGAGCATTGAAGAGATCAAC
GCCCTGGAAGAAGAAGATAAAAACAGCGGTAAAGCACCGCGCGCCCAGTATGTACT
GGCGGAGCAGGTGACTCGTCTGGTTCACGGTGAAGAAGGTTTACAGGCGGCAAAAC
GTATTACCGAATGCCTGTTCAGCGGTTCTTTGAGTGCGCTGAGTGAAGCGGACTTCG
AACAGCTGGCGCAGGACGGCGTACCGATGGTTGAGATGGAAAAGGGCGCAGACCTG
ATGCAGGCACTGGTCGATTCTGAACTGCAACCTTCCCGTGGTCAGGCACGTAAAACT
ATCGCCTCCAATGCCATCACCATTAACGGTGAAAAACAGTCCGATCCTGAATACTTC
TTTAAAGAAGAAGATCGTCTGTTTGGTCGTTTTACCTTACTGCGTCGCGGTAAAAAG
AATTACTGTCTGATTTGCTGGAAAtaa

pB1U Ec-OMeY-RS
Same as Ec-BPA-RS but with the following active site mutations:
Y37V, D182S, F183M, L186A

pB1U H2-BPA-RS
Same as above but with the following aaRS sequence:
atgGCAAGCAGTAACTTGATTAAACAATTGCAAGAGCGGGGGCTGGTAGCCCAGGTG
ACGGACGAGGAAGCGTTAGCAGAGCGACTGGCGCAAGGCCCGATCGCGCTCGGGTG
CGGCTTCGATCCTACCGCTGACAGCTTGCATTTGGGGCATCTTGTTCCATTGTTATGC
CTGAAACGCTTCCAGCAGGCGGGTCACCGTCCGATCGCGCTGGTTGGTGGTGCGACC
GGTCTGATTGGCGACCCGAGCGGCAAGAAAAGCGAGCGTACCCTGAACGCGAAGG
AAACCGTTGAAGCGTGGAGCGCGCGTATCAAAGAACAGCTGGGTCGTTTCCTGGAC
TTTGAGGCGGATGGCAACCCGGCGAAGATTAAAAACAACTATGACTGGATCGGTCC
GCTGGATGTGATTACCTTCCTGCGTGATGTGGGCAAGCACTTTAGCGTTAACTACAT
GATGGCGAAAGAGAGCGTTCAGAGCCGTATCGAAACCGGTATTAGCTTCACCGAGT
TTAGCTACATGATGCTGCAAGCGTATGGTTTCCTGCGTGCCTACGAAACCGAAGGCT
GCCGTCTGCAGATCGGTGGCAGCGATCAATGGGGTAACATCACCGCGGGCCTGGAA
CTGATTCGTAAGACCAAAGGTGAAGCGCGTGCGTTTGGCCTGACCATCCCGCTGGTG
ACCAAGGCGGACGGTACCAAGTTTGGCAAAACCGAAAGCGGTACCATTTGGCTGGA
TAAGGAGAAAACCAGCCCGTACGAATTCTATCAGTTTTGGATCAACACCGACGATC
GTGACGTTATTCGTTACCTGAAGTATTTCACCTTTCTGAGCAAAGAGGAAATCGAAG
CGCTGGAGCAGGAACTGCGTGAGGCGCCGGAAAAGCGTGCGGCGCAAAAAGCGCT
GGCGGAGGAAGTGACCAAACTGGTTCACGGTGAGGAAGCGCTGCGTCAGGCGATCC
GTATTAGCGAAGCGCTGTTTAGCGGTGATATCGCGAACCTGACCGCGGCGGAGATT
GAACAAGGCTTCAAGGACGTGCCGAGCTTTGTTCACGAAGGTGGCGATGTGCCGCT
GGTTGAGCTGCTGGTTAGCGCGGGTATCAGCCCGAGCAAACGTCAGGCGCGTGAAG
ACATCCAAAACGGTGCGATTTACGTGAACGGCGAGCGTCTGCAAGATGTTGGCGCG

ATTCTGACCGCGGAACACCGTCTGGAAGGTCGTTTTACCGTTATCCGTCGTGGCAAG
AAGAAATACTATCTGATTCGTTATGCGTAA

pB1U H2-OMeY-RS
Same as H2-BPA-RS but with the following active site mutations:
Y37V, D176S, F180M, L183A

pB1U H6 BPA-RS
Same as above but with the following aaRS sequence:
atgGCAAGCAGTAACTTGATTAAACAATTGCAAGAGCGGGGGCTGGTAGCCCAGGTG
ACGGACGAGGAAGCGTTAGCAGAGCGACTGGCGCAAGGCCCGATCGCGCTCGGGTG
CGGCTTCGATCCTACCGCTGACAGCTTGCATTTGGGGCATCTTGTTCCATTGTTATGC
CTGAAACGCTTCCAGCAGGCGGGCCACAAGCCGGTTGCGCTGGTAGGCGGCGCGAC
GGGTCTGATTGGCGACCCGAGTTTCAAAGCTGCCGAGCGTAAGCTGAACACCGAAG
AAACTGTTCAGGAGTGGGTGGACAAAATCCGTAAGCAGGTTGCCCCGTTCCTCGATT
TCGACTGTGGAGAAAACTCTGCTATCGCGGCGAACAACTATGACTGGTTCGGCAATA
TGAATGTGCTGACCTTCCTGCGCGATATTGGCAAACACTTCTCCGTTAACCAGATGA
TCAACAAAGAAGCGGTTAAGCAGCGTCTCAACCGTGAAGATCAGGGGATTTCGTTC
ACTGAGTTTTCCTACAGCCTGTTGCAGGGTTATGGGTTCGCCTGTGCTAACAAACAG
TACGGTGTGGTGCTGCAAATTGGTGGTTCTGACCAGTGGGGTAACATCACTTCTGGT
ATCGACCTGACCCGTCGTCTGCATCAGAATCAGGTGTTTGGCCTGACCGTTCCGCTG
ATCACTAAAGCAGATGGCACCAAATTTGGCAAAACCGAAAGCGGTACCATTTGGCT
GGATAAGGAGAAAACCAGCCCGTACGAATTCTATCAGTTTTGGATCAACACCGACG
ATCGTGACGTTATTCGTTACCTGAAGTATTTCACCTTTCTGAGCAAAGAGGAAATCG
AAGCGCTGGAGCAGGAACTGCGTGAGGCGCCGGAAAAGCGTGCGGCGCAAAAAGC
GCTGGCGGAGGAAGTGACCAAACTGGTTCACGGTGAGGAAGCGCTGCGTCAGGCGA
TCCGTATTAGCGAAGCGCTGTTTAGCGGTGATATCGCGAACCTGACCGCGGCGGAG
ATTGAACAAGGCTTCAAGGACGTGCCGAGCTTTGTTCACGAAGGTGGCGATGTGCC
GCTGGTTGAGCTGCTGGTTAGCGCGGGTATCAGCCCGAGCAAACGTCAGGCGCGTG
AAGACATCCAAAACGGTGCGATTTACGTGAACGGCGAGCGTCTGCAAGATGTTGGC
GCGATTCTGACCGCGGAACACCGTCTGGAAGGTCGTTTTACCGTTATCCGTCGTGGC
AAGAAGAAATACTATCTGATTCGTTATGCGTAA
